# The effect of semantic content on the perception of audiovisual movieclips

**DOI:** 10.1101/2024.01.24.576956

**Authors:** Anastasia Maria Kesoglou, Kyriaki Mikellidou

**Affiliations:** Open University of Cyprus; University of Limassol

**Keywords:** Crossmodal integration, Audiovisual integration, Semantic congruency, Semantic audiovisual movies, Multimodal perception, Verbs, Nouns

## Abstract

Our brain is skilled with the ability to perceive and process multimodal stimuli. This process known as crossmodal perceptual integration, has been in the research spotlight for a long time, providing evidence for the integration of information coming from different modalities. Prior research mostly utilized pictures and focused on the semantic content of a single sound or word. The present study aims to investigate crossmodal perceptual integration in realistic conditions using short movieclips (1500ms) and auditory meaningful three-word sentences to evaluate target detection in terms of accuracy and response times. Two experimental tasks were developed using PsychoPy, where participants had to indicate whether a target (noun for Experiment 1, verb for Experiment 2) was present or absent. In trials without a target, target-related information was always present, either through one of the two senses (vision or audition; incongruent condition) or through both senses (congruent condition). We observed superior performance when the target was absent generally (M_exp1_ =93.9%, SD_exp1_ =0.04, M_exp2_ =84.2%, SD_exp2_ =0.182) compared to when it was present (M_exp1_ =82.3%, SD_exp1_ =0.143, M_exp2_ =73.7%, SD_exp2_ =0.184). Moreover, superior performance was noted in incongruent target-related movieclips, which significantly decreased during congruent target-related movieclips. In Experiment 1, we observed that in the audio condition when the target-related word was a noun, participant performance was superior compared to when it was a verb (M=99.4% vs. M=86.7%; t_incVerb vs. incNoun_ =-8.428, p=.001). In Experiment 2, the judgment scores were similarly high in incongruent movieclips and significantly lower in congruent ones regardless of whether the target-related information presented was a verb or noun. The present results provide evidence regarding the role of complexity of semantics, and especially the diverse role verbs and nouns could play in crossmodal perceptual integration in more realistic situations. Our findings can enrich the content of learning techniques, as well as the design of AI models, by taking advantage of the supporting role of semantic audiovisual information, while taking into consideration the potential confusion that the complexity of semantic information can induce to perception experience.

## Introduction

We perceive the world with our five senses: vision, audition, taste, olfaction, and touch -which includes tactile and temperature sense, as well as pressure. The process referring to the temporal and spatial binding of two or more perceptual features rooting from the same or different sensory modalities refers to what is known as crossmodal integration (Lalanne & Lorenceau, 2004; Lachs, 2023). Examples of how multimodal interactions work and of their outcomes come from a range of experimental settings utilizing all possible crossmodal and multimodal stimulus pairings. For instance, many researchers focused on the level of enhancement of one modality over the other during a crossmodal perceptual task, where crossmodal signals were either congruent (temporally/spatially) or incongruent. (Stein et al., 1996; Shams, Kamitani, & Shimojo, 2000; Calvert, Campbell, & Brammer, 2000; Vroomen & de Gelder, 2000; Morein-Zamir, Soto-Faraco & Kingstone, 2003; Laurienti et al., 2004; Ro et al, 2004; Sakai., 2005; Zampini et al., 2005; Chen & Spence, 2010; van de Groen et al., 2013; Cox & Hong, 2015; Li et al., 2019; Brandman et al., 2020; Rekow et al., 2022; Woods et al., 2023). On the other hand, a suppression effect of sensory cues contributes to a class of perceptual illusions (Tsuchiya, 2008). According to Bruns (2019), these perceptual illusions arise when, for example, the highest weight is being given on visual cues, influencing in this way, the perception of an audio cue (i.e., its location). Perceptual illusions have been mostly reported in settings that include the visual modality (Violentyev, Shimojo, & Shams; 2005; Kammers et al., 2009; Moscatelli et al. 2015; Kang, Sah, & Lee, 2021). In such, evidence has been provided showing that auditory information can qualitatively and quantitatively alter the perception of a visual stimulus. The visual illusion of perceiving multiple flashes because of the presence of multiple auditory beeps is an example of qualitative change of visual input caused by an audio-cue (Shams, Kamitani, & Shimojo, 2000). Respectively, the influence of sound on perception of light intensity applies as an example of quantitative change (Stein et al., 1996).

### The Role of Semantics

The presence of multiple crossmodal cues can enhance behavioral performance by speeding responses, increasing accuracy, and improving stimulus detection (Laurienti et al., 2004). Studies focusing on brain activation patterns during perception of semantically congruent and incongruent audiovisual stimuli suggest that there is a strong relation between congruent audiovisual stimuli and facilitation of neural representations of semantic categories or concepts (Li et al., 2011). Thus, it is of great interest to examine the role of semantic content in the perception of audiovisual stimuli. As Laurienti et al. suggested in 2004, by studying the importance of semantic content on crossmodal effects, the research community could offer more insights into neural operations and constituents of crossmodal information processing. There have been a few but very interesting studies which focused on the impact of semantic sounds on the perception of visual stimuli (i.e., static pictures) and the semantic congruency between crossmodal stimuli. Chen and Spence (2010) conducted a series of experiments to assess the effect of audiovisual semantic congruency specified on the identification of masked visual targets. The results indicate a shared semantic system in which neural representations of crossmodal stimuli interact. This crossmodal semantic interaction depends on a short-term buffer responsible for handling semantic representations. In addition, Fujisaki et al. (2014) found strong associations in material perception (e.g., glass, plastic, ceramic, paper) of simultaneous audiovisual stimuli. In particular, participants’ material categorization of an object shown on video was influenced by the material sound (e.g., the sound of a vegetable’s surface when it was hit by a wooden mallet or the sound of glass when hit by a wooden mallet, etc.). Their results indicated that in irrelevant audiovisual signals the perception of the material was modified depending on the combination of both the audio sound and the visual clip (for example the sound of vegetable hit by a wooden mallet and the visual clip of a glass were combined and perceived as a video that shows a plastic bottle).

The significant impact of semantics on audiovisual integration in perception has been also examined in the case of bistable figures. In such, Hsiao et al. (2012) indicate that background auditory soundtracks such as the voice of a young or old female can alter the predominant perception of a bistable figure such as a “wife” or “mother in-law” figure. This crossmodal semantic effect occurred in respect to manipulation of visual fixation and showed to interact with voluntary attention. Recently, Williams et al. (2022) used a psychophysical task consisting of pairs of naturalistic sounds and noisy visual images. In their first experiment an ambiguous visual image which referred to two distinct objects (identical shadows of two different objects) switched from obscured to clear view while a naturalistic sound played. The incidental sound was either coherent with one of the two objects or completely irrelevant. Participants were asked to recreate the target morph they previously saw as accurately as possible using a continuous report line and press the button when they were ready to submit their answer. No sound was playing during the response phase. In the second experiment, sounds were played during the continuous report phase whereas in the second part of the experiment half of all blocks had no sound. A further experiment was conducted to examine potential high-level semantic representations implicated by sounds, presenting the full length of the naturalistic sound followed by the visual morph after a 3-s delay. The authors observed a continuous integration of temporally and semantically congruent audiovisual inputs and explained the perception of visual objects as a function of naturalistic auditory context, since the latter provides and enriches visual perception with complementary information, which is both independent and diagnostic (Williams et al., 2022).

Except for audiovisual simple signals or sounds, research approaches have also examined the perceptual integration of linguistic unimodal and crossmodal information. In the case of unimodal stimuli, there is evidence regarding the influence of language on a specific visual attribute when the content of the presented written sentence and visual attribute are semantically congruent (Pelekanos & Moutoussis, 2011). In the case of crossmodal stimuli, semantic congruency effects of auditory stimuli on the processing of visual cues were examined according to two factors: synchronization and categorical specificity. In such, Chen and Spence (2018) used either naturalistic sounds or spoken words in combination with pictures or printed words. Seven stimulus onset asynchronous (SOA) conditions were utilized in the experiments, and the task given to the participants required speeded categorization judgements to whether each cue presented belonged to the living or nonliving objects category. Both congruency and inhibitory effects were reported for different SOA conditions.

In respect with the unity assumption theory –according to which modulation of multisensory integration is the result of one observer’s assumption or beliefs that multiple unisensory signals root from the same source (Chen & Spence, 2017), Thomas and Shiffrar (2013) studied the circumstances under which the temporal relationship between auditory and visual stimuli modulate this inhibitory effect in perception. In their first experiment, participants were asked to identify covered-up point-light walkers while listening to footsteps that were either synchronous or out-of-phase with the point-light footfalls. In the second experiment the rhythm factor was added in the auditory and visual streams, so this time, participants were again asked to detect point-light walkers but the auditory cues where either footsteps or tone sounds, synchronous with the point-light footfalls or temporally random (completely decoupled from the motion). The findings suggest that in all conditions, relative timing of auditory and visual stimuli was not a critical factor for enhanced visual sensitivity in detecting actions, but semantic congruency was (Thomas & Shiffrar, 2013). Recently, Uno & Yokosawa (2022) took into consideration the spatiotemporal congruency effects between modalities, and examined their potential relation with temporal recalibration of audiovisual stimuli during perceptual integration processing. For this purpose, they conducted simultaneity judgment tasks consisting of audiovisual pairs (audio pitch either high or low, and visual cycle either presented above fixation point or below) that were synchronous or asynchronous. In each block, participants were initially completing adaptation trials for 60s where alternating auditory and visual stimuli were presented with no time interval differences. Their results showed selective recalibration of asynchronous but semantically congruent audiovisual signals.

### Purpose

The aim of the current study is to introduce more realistic aspects in the examination of crossmodal integration with the simultaneous presentation of short movieclips and auditory three-word sentences. Prior experiments utilized mostly stationary visual cues and were limited in the semantic content of a naturalistic sound or a single word. The main hypothesis refers to whether and to what extent audiovisual semantic information affects speed and accuracy in judgments in detecting the presence or absence of a target. Complementary to this, the present study is the first to introduce target absent trials which always contain target-related information either presented through vision, audition, or both senses. In such trials, audio sentences include a semantic target-related noun or verb. To date no other study investigated the potential influence of the type of semantics (verb vs. noun) on the perception of audiovisual stimuli, especially of movieclips. If target-related information is considered useful for the neural system to identify the absence of the target then we expect that performance (both in terms of speed and accuracy) would be improved, otherwise if it is considered noise, we expect to observe a compromised performance.

## Materials and Methods

Experimental procedures were approved by the Cyprus National Bioethics Committee (ΕΕΒΚ ΕΠ 2018.01.183) and are in line with the Declaration of Helsinki abiding to ethical standards that promote and ensure respect for all human participants, while protecting their health and rights. Research practices conformed to generally accepted scientific principles and were all based on a thorough knowledge of relevant scientific literature.

### Experiment 1

### Participants

We tested a total of 38 participants, but data from two participants was excluded as one requested not to be included in the data analysis after completing the task, and the other reported that they faced technical issues during the experiment (24 female; age range 18–60 with three missing values; mean age = 30). Inclusion criteria involved normal or corrected-to-normal vision, no neurological disorders, no learning disabilities. Participants were mainly recruited from undergraduate and postgraduate classes at the University of Cyprus and the University of Limassol receiving course credit in exchange for their participation. Each participant was provided written detailed instructions regarding the experimental online task, and the equipment that was required in order to run the task.

### Experimental Design

### Stimuli

#### Targets

In Experiment 1 the target was always a noun. There were 12 targets, each one presented in each trial randomly, through both senses: ***(a)*** knife, ***(b)*** lemons, ***(c)*** (wedding) ring, ***(d)*** children, ***(e)*** towel, ***(f)*** apron, ***(g)*** bus, ***(h)*** (bus) stop, ***(i)*** stairs, ***(j)*** uphill, ***(k)*** hairdryer, ***(l)*** hair salon. In the visual modality a line-drawing of the target was presented, while in the audio modality the target-word was verbalized.

#### Visual clips

There were 12 short scenes of 1500 ms cut out from the short movie “37 Days” directed by Nikoleta Leousi (from which we took written permission of use), standardized on familiarity and complexity. Standardized on familiarity refers to the scenes since they were all coming from real-word routines like walking and cooking, while standardized on complexity means that all videos included at least one and not more than two movements in the scene presented. As shown in the *Table 1* (see Appendix) these were: ***(a)*** a hand stirring lemon juice in a glass with lemon slices on the foreground, ***(b)*** a hand cutting an onion, ***(c)*** pregnant woman caressing her belly, ***(d)*** children with parents at a waiting hall, ***(e)*** a woman and an old man sitting in a bus, ***(f)*** a pregnant woman arriving at a crowded bus stop, ***(g)*** a woman going up the stairs, ***(h)*** a woman walking on a hill, ***(i)*** two hairdressers with the one taking of a towel from a woman in a hair salon, ***(j)*** a sitting man wiping a telephone on his apron, ***(k)*** a pregnant woman staring at a mirror and a hairdryer, ***(l)*** a pregnant woman arriving at a hair salon. These were presented in full-screen mode on a gray background. The size of the screen varied according to the size of each participant’s screen.

**Table 1a.**
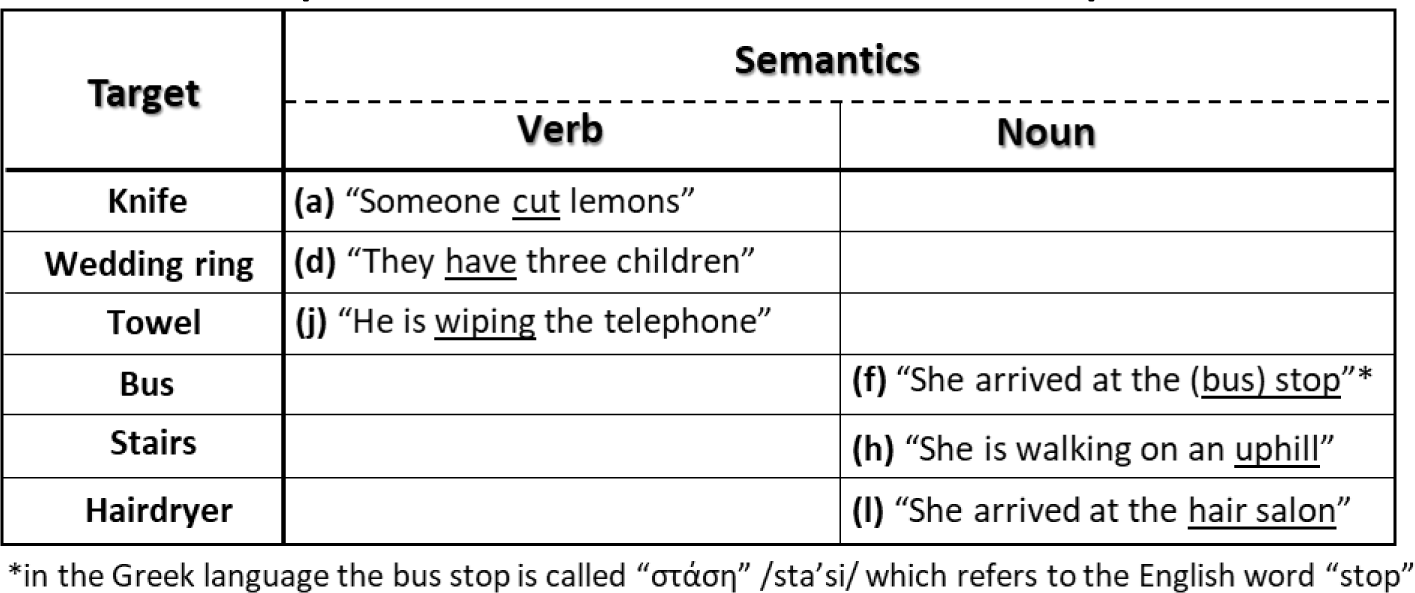
The experimental conditions for semantics in Experiment 1.

**Table 1b.** The experimental conditions for semantics in Experiment 2.

| Target | Semantics |  |
| --- | --- | --- |
|  | Verb | Noun |
| Sit |  | (a) "She is climbing on a <u>chair</u>" |
| Drink |  | (h) "He is stirring a <u>juice</u>" |
| Fold |  | (l) "She is taking off the <u>apron</u>" |
| Run | (c) "She is <u>walking</u> in the neighborhood" |  |
| Caress | (f) "She is <u>hugging</u> a friend" |  |
| Clean | (i) "She is <u>wiping</u> the hair" |  |

#### Audio clips

There were 12 semantic audio sentences in the Greek language consisting of three-word sentences, spoken by an AI voice using the mobile application “Text To Speech” developed by STCodesApp and provided by Google Commerce Ltd (pitch and volume were defined at 50%, and speed at 26%). In particular, the 12 sentences (translated in English) are: ***(a)*** “Someone <u>cut</u> lemons”; ***(b)*** “He is using the knife”; ***(c)*** “She is wearing a wedding ring”; ***(d)*** “They <u>have</u> three children”; ***(e)*** “They are sitting in the bus”; ***(f)*** “She arrived at the <u>bus stop</u>”; ***(g)*** “She is going up the stairs”; ***(h)*** “She is walking <u>uphill</u>”; ***(i)*** “She removes the towel; ***(j)*** “He is <u>wiping</u> the telephone”; ***(k)*** “She found the hairdryer”; ***(l)*** “She arrived at the <u>hair salon</u>”. As shown in *Table 2a*, the sentences belonged in one of two basic semantic categories according to the type of the word (verb vs. noun) which was semantically related to the target expected in the movieclip. The category *verb* included the sentences ***a*** for the target “knife”, ***d*** for the target “ring”, ***j*** for the target “towel”, and the category *noun* included the sentences ***f*** for the target “bus”, ***h*** for the target “stairs”, ***l*** for the target “hairdryer”. The duration of all sentences was identical to the duration of the short movieclips (i.e.1500 ms).

**Table 2.**
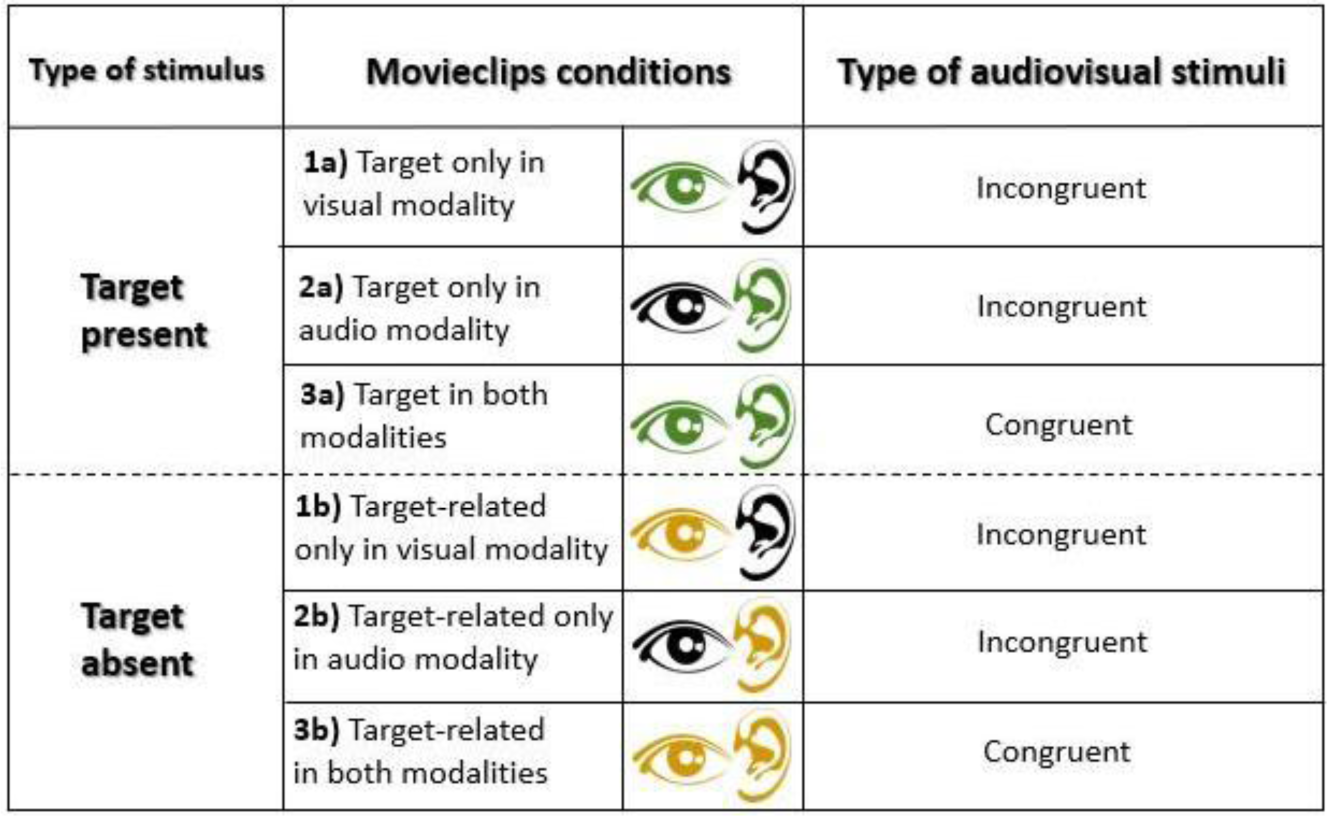
The experimental conditions. The experimental conditions were distinguished in target present and target absent trials. Movieclips in target present trials were distinguished according to the modality in which the target stimulus was presented: 1a, 2a, 3a. In target absent trials movieclips were distinguished according to the modality in which the target-related information was presented: 1b, 2b, 3b. When the target or target-related information was presented only in one of the two modalities, the movieclips were characterized as incongruent, while when it was presented in both, the movieclips were characterized as congruent.

| Type of stimulus | Movieclips conditions |  | Type of audiovisual stimuli |
| --- | --- | --- | --- |
| **Target present** | **1a)** Target only in visual modality |  | Incongruent |
|  | **2a)** Target only in audio modality |  | Incongruent |
|  | **3a)** Target in both modalities |  | Congruent |
| **Target absent** | **1b)** Target-related only in visual modality |  | Incongruent |
|  | **2b)** Target-related only in audio modality |  | Incongruent |
|  | **3b)** Target-related in both modalities |  | Congruent |

### Procedure

Initially participants were debriefed, and were given the information to ask questions regarding the task. They were also reminded to remove any distracting item, to run the task in a quiet place alone, and to ensure their headphones were plugged in, if using any. When participants were ready, the URL was provided to them in order to start the experimental task. Participants were first asked to provide their age, gender, and country; then detailed instructions about the task were provided through an AI voice and pictures, and finally an example trial was provided. As soon as participants were ready to start the main trials, they responded by pressing the right arrow on their keyboard. The experimental timeline is shown in detail in *Figure 1*. A fixation point (400 ms) was firstly presented, followed by a line drawing of the target stimulus in the movieclip. This was presented simultaneously with a written and a verbal label (1000 ms). Another fixation point (500 ms) followed, and the movieclip (1500 ms) was presented right after. Finally, a line drawing of the two possible response key-arrows appeared accompanied by the label “Did you see and/or hear the target?”. Participants were instructed to answer as quickly and accurately as possible whether they perceived the target in the movieclip (through either vision/audition or both).

**Figure 1.**
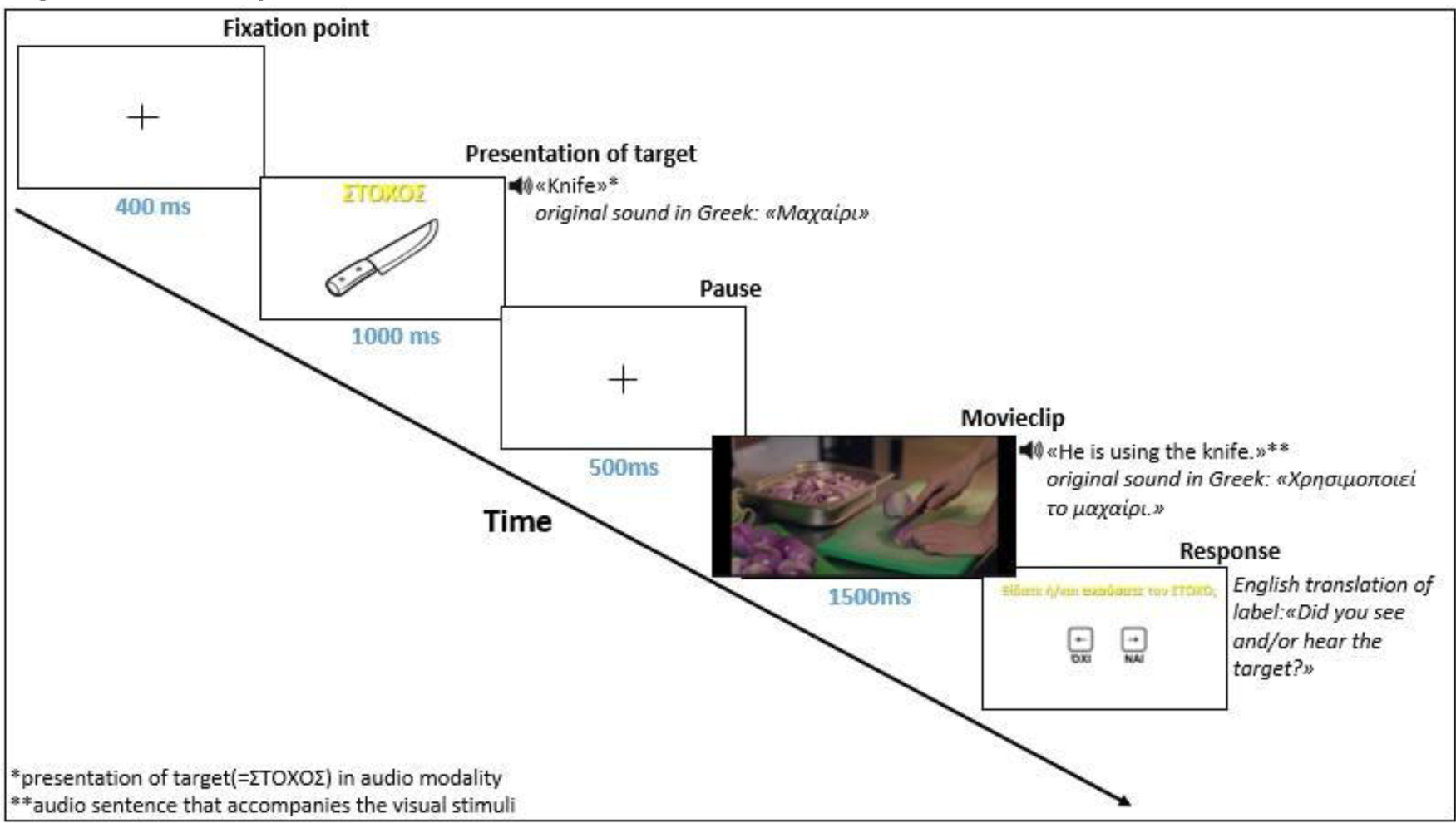
The experimental timeline. The example given here corresponds to a trial in which both the audio and visual stimuli included the target *Knife* (congruent movieclip). After every movieclip the picture with the response keys (right arrow for “YES” and left arrow for “NO”) appeared immediately. Response recording and reaction times (RT) started right from the moment this picture appeared (RT = inter-trial interval (ITI)).

The task was self-paced, producing a variable inter-trial interval (ITI) which was identical to the participant’s RT for that specific trial. Then, a new trial started immediately. There were no breaks available since the task could not be paused. However, each participant was informed that the task can be terminated at any time by pressing the escape button on their keyboard if they did not wish to proceed with the experiment.

### Experimental Conditions

The experimental task was designed using the PsychoPy software (Peirce et al., 2019) and was online running through pavlovia.org. Accuracy in responses and RT for each trial were recorded by the built-in keyboard backend PsychToolbox -Psychophysics Toolbox extensions (Kleiner et al., 2007), for data collection of keyboard input in PsychoPy. The experiment was based on two target conditions; target present and target absent, where the target absent condition included only trials with at least one modality presenting information semantically related to the target (thus, we refer to these as target-related stimuli). In addition, the target and target-related stimuli were equiprobably presented through modalities as follows: ***i)*** target presented only in the visual modality, ***ii)*** target presented only in the audio modality, ***iii)*** target presented in both audio and visual modalities, ***iv)*** target-related only in the visual modality, ***v)*** target-related only in the audio modality, ***vi)*** target-related in both audio and visual modalities. By applying the types of stimulus (target and target-related) in the above six possible conditions of modality, we received the following audiovisual combinations: ***(I)*** target/target-related stimulus only presented in an audio clip accompanied by an incongruent visual clip; ***(II)*** target/target-related only presented in a visual clip accompanied by an incongruent audio clip; ***(III)*** target/target-related stimulus presented in both modalities. The first two (I and II) combinations have been grouped under the term “incongruent audiovisual stimuli” whereas the latter under the term “congruent audiovisual stimuli” (see *Table 3*). Thus, an example for incongruent audiovisual movieclip with the target stimulus in: ***1a)*** *visual modality* is a visual clip including the target “knife” accompanied by an audio sentence that does not have the target-word “knife”; ***2a)*** *audio modality* is an audio sentence including the target-word “knife” accompanied by a visual clip that does not include the target “knife”; whereas an example for congruent audiovisual movieclip with the target stimulus in ***3a)*** *both visual and audio modalities* is a visual clip that includes the target “knife” accompanied by an audio sentence including the target-word “knife”. Respectively, an example for incongruent audiovisual movieclip with the target-related stimulus in: ***1b)*** *visual modality* is a target-related visual clip for the target “knife” (i.e., a hand stirring lemon juice in a glass with lemon slices on the foreground) accompanied by an audio sentence that does not have the target-word “knife” or a target-related word to it, ***2b)*** *audio modality* is a target-related audio sentence for the target-word “knife” accompanied by a visual clip that does not include the target “knife” or a target-related object or action to it; whereas an example for congruent audiovisual movieclip with the target-related stimulus in ***3b)*** *both visual and audio modalities* is a target-related visual clip for the target “knife” accompanied by the target-related audio sentence for the target-word “knife”.

**Table 3.**
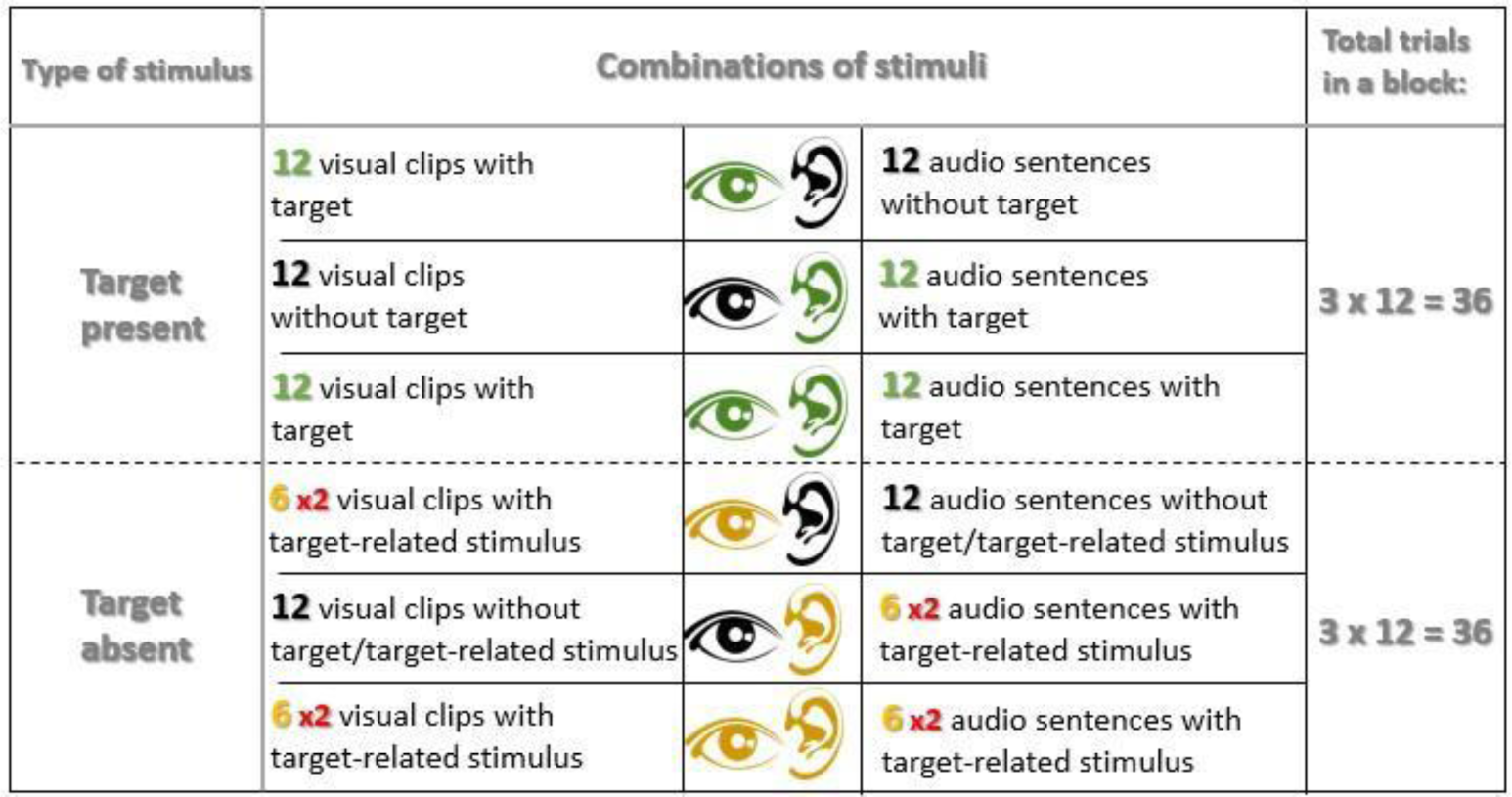
Total number of movieclips. Movieclips in target present trials were built out of the combinations of target and target absent audio and visual stimuli, following the rule that at least one modality includes the target. In target absent trials, movieclips were the outcome of the combinations of target-related and target absent stimuli, following the rule that at least one modality includes target-related information. In target absent trials, we replaced the targets with related information only for six targets. Therefore, the total target-related clips used for the combinations in each modality condition was six.

| Type of stimulus | Combinations of stimuli |  |  | Total trials in a block: |
| --- | --- | --- | --- | --- |
| **Target present** | 12 visual clips with target |  | 12 audio sentences without target | **3 x 12 = 36** |
|  | 12 visual clips without target |  | 12 audio sentences with target |  |
|  | 12 visual clips with target |  | 12 audio sentences with target |  |
| **Target absent** | 6 x 2 visual clips with target-related stimulus |  | 12 audio sentences without target/target-related stimulus | **3 x 12 = 36** |
|  | 12 visual clips without target/target-related stimulus |  | 6 x 2 audio sentences with target-related stimulus |  |
|  | 6 x 2 visual clips with target-related stimulus |  | 6 x 2 audio sentences with target-related stimulus |  |

Target trials and target absent trials occurred the same number of times. The number of stimuli presented during the task was for the **A)** target present condition: six audio sentences, six visual clips, with 36 pairings of the visual and audio stimuli. However, for **B)** target absent condition (in which target-related information was present) there were again six audio sentences, six visual clips, but with 30 pairings plus the repetition of the six congruent audiovisual pairings (see *Table 4*). The experimental task was running in four repetitions (4 blocks). Therefore, the total number of trials was 288 (144 target trials and 144 target-related trials), which were equally distributed for each audiovisual target and target-related condition: audio modality (48 trials with target stimuli; 48 with target-related stimuli); visual modality (48 trials with target stimuli and 48 with target-related stimuli); audiovisual modality (48 trials with target stimuli and 48 with target-related stimuli). For the target-related trials, audio sentences were also equally distinguished with respect to the category of semantics they contained. That is, whether the word which was semantically related to the expected target, was a verb or a noun (24 audio sentences including a semantically related verb and 24 including a semantically related noun).

**Table 4a.** Post Hoc Comparisons for Mean Proportion Correct in target present and target absent conditions (Modality * Target presence) in Experiment 1.

|  |  | Experiment 1 |  |  |  |
| --- | --- | --- | --- | --- | --- |
|  |  | Mean Difference | SE | t | P<sub>Holm</sub> |
| Audio, yes | Visual, yes | -0.193 | 0.037 | -5.275 | < .001 |
|  | Audiovisual, yes | -0.333 | 0.037 | -9.094 | < .001 |
|  | Audio, no | -0.284 | 0.040 | -7.044 | < .001 |
| Visual, yes | Audiovisual, yes | -0.140 | 0.037 | -3.819 | 0.002 |
|  | Visual, no | -0.146 | 0.040 | -3.619 | 0.004 |
| Audiovisual, yes | Audiovisual, no | 0.083 | 0.040 | 2.051 | 0.257 |
| Audio, no | Visual, no | -0.055 | 0.037 | -1.512 | 0.553 |
|  | Audiovisual, no | 0.033 | 0.037 | 0.899 | 0.741 |
| Visual, no | Audiovisual, no | 0.088 | 0.037 | 2.411 | 0.121 |

**Table 4b.** Post Hoc Comparisons for Mean Proportion Correct in target present and target absent conditions (Modality * Target presence) in Experiment 2.

|  |  | Experiment 2 |  |  |  |
| --- | --- | --- | --- | --- | --- |
|  |  | Mean Difference | SE | t | P<sub>Holm</sub> |
| Audio, yes | Visual, yes | -0.229 | 0.046 | -5.000 | < .001 |
|  | Audiovisual, yes | -0.466 | 0.046 | -10.180 | < .001 |
|  | Audio, no | -0.436 | 0.057 | -7.689 | < .001 |
| Visual, yes | Audiovisual, yes | -0.237 | 0.046 | -5.180 | < .001 |
|  | Visual, no | -0.182 | 0.057 | -3.219 | 0.009 |
| Audiovisual, yes | Audiovisual, no | 0.302 | 0.057 | 5.335 | < .001 |
| Audio, no | Visual, no | 0.024 | 0.046 | 0.532 | 0.948 |
|  | Audiovisual, no | 0.272 | 0.046 | 5.940 | < .001 |
| Visual, no | Audiovisual, no | 0.248 | 0.046 | 5.408 | < .001 |

### Experiment 2

### Participants

Thirty-four volunteers participated (mean age 30, age range 22-40 with five missing values; 20 female and 14 male). All other information is identical to Experiment 1.

### Experimental Design

The procedure and experimental conditions were the same with Experiment 1, but in Experiment 2 the target stimulus was always a verb.

### Stimuli

#### Targets

There were six verb targets, each one presented in each trial randomly, through both senses: ***(a)*** climb on, ***(b)*** sit, ***(c)*** walk, ***(d)*** run, ***(e)*** hug, ***(f)*** caress, ***(g)*** drink, ***(h)*** stire, ***(i)*** wipe, ***(j)*** clean, ***(k)*** fold, ***(l)*** take off. In the visual modality a line-drawing of the target was presented, while in the audio modality the target-word was verbalized.

#### Visual clips

Similar to Experiment 1, there were 6 short scenes of 1500 ms cut out from the short movie “37 Days”. These videos were: ***(a)*** a woman in her kitchen climbing on a chair, ***(b)*** a woman sitting on a chair with a man staring at her on the background, ***(c)*** a man running, ***(d)*** a pregnant woman walking, ***(e)*** a pregnant woman caressing her belly, ***(f)*** two women hugging, ***(g)*** a sitting pregnant woman drinking a juice and a man sitting beside her, ***(h)*** someone stirring a lemon juice, ***(i)*** two hairdressers with the one wiping the hair of the costumer using a towel, ***(j)*** a sitting man cleaning potatoes, ***(k)*** a pregnant woman folding clothes, ***(l)*** a pregnant woman taking off her apron. They were presented in full-screen mode on a gray background. All other information was identical to Experiment 1.

#### Audio clips

Similar to Experiment 1, there were 12 semantic audio sentences. In particular, the 12 sentences (as translated in English) are: ***(a)*** “She is climbing on a <u>chair</u>”; ***(b)*** “She is sitting to have a break”; ***(c)*** “She is <u>walking</u> in the neighborhood”; ***(d)*** “He is running on the pave way”; ***(e)*** “She is caressing the belly”; ***(f)*** “She is <u>hugging</u> a friend”; ***(g)*** “She is drinking a lemonade”; ***(h)*** “He is stirring a <u>juice</u>”; ***(i)*** “She is <u>wiping</u> the hair”; ***(j)*** “He is cleaning the potatoes”; ***(k)*** “She is folding the clothes”; ***(l)*** “She is taking off the <u>apron</u>”. As shown in *Table 2b*, the sentences were distinguished in two basic semantic categories according to the type of the word (verb vs. noun) which was semantically related to the target expected in the movieclip. The category *verb* included the sentences ***c*** for the target “run”, ***f*** for the target “caress”, ***i*** for the target “clean”, and the category *noun* included the sentences ***a*** for the target “sit”, ***h*** for the target “drink”, ***l*** for the target “fold”. All other information was identical to Experiment 1.

### Data Analysis

In both Experiments, data for each participant was collected and automatically saved in excel files from the Pavlovia platform. For individual participants, mean proportion correct and mean reaction times (RT) were calculated (for each condition) using JASP software version 0.17.1. To test for an effect of target presence on proportion correct and RT, we performed two separate two-way repeated measures analyses of variance (ANOVA), with Modality (three levels: audio, visual, audiovisual) and Target presence (two levels: target present, target absent) as the main factors.

We also tested whether the type of a semantically related word (verb or noun) in a three-word sentence affects accuracy and reaction times when the target is absent in both modalities. Thus, another two-way repeated measures ANOVA was conducted with Congruency (congruent vs. incongruent) and Semantics (verb vs. noun) as main factors. In this way, we tested for potential differences in results (proportion correct of responses and RTs) between congruent audiovisual clips when containing a target-related verb vs. noun, as well as between the incongruent and the congruent audiovisual clips with a semantically related verb, and noun respectively.

Moreover, for all ANOVA, Greenhouse-Geisser correction was applied when the assumption of sphericity was violated.

For the analysis, outlier trials that were 3 standard deviations above or below each participant’s mean RT were removed. All trials (correctly and erroneously answered) were used for the analysis of RTs presented in the main text. In the Appendix, the RT analysis of only correctly answered trials can also be found (Appendix: *Figures A3*, *A4*).

## Results

### Mean Proportion Correct

### Experiment 1: Target is always a noun

The results indicate a statistically significant difference in mean proportion correct responses depending on the presence of the target and the modality in which the target or target-related information was presented. *Figure 3a-b* shows results from movieclips with incongruent information where the target noun is presented only in the auditory or visual modality. We observed a similar pattern of results: accuracy in judgments was low when the target was present only in the audio (M_audio_= 0.647, SD_audio_= 0.305) and only in the visual modality (M_visual_=0.841, SD_visual_= 0.235), and increased when the target was absent from both modalities (M_audio_= 0.931, SD_audio_= 0.038; M_visual_= 0.986, SD_visual_= 0.018) while variance was reduced.

**Figure 3.**
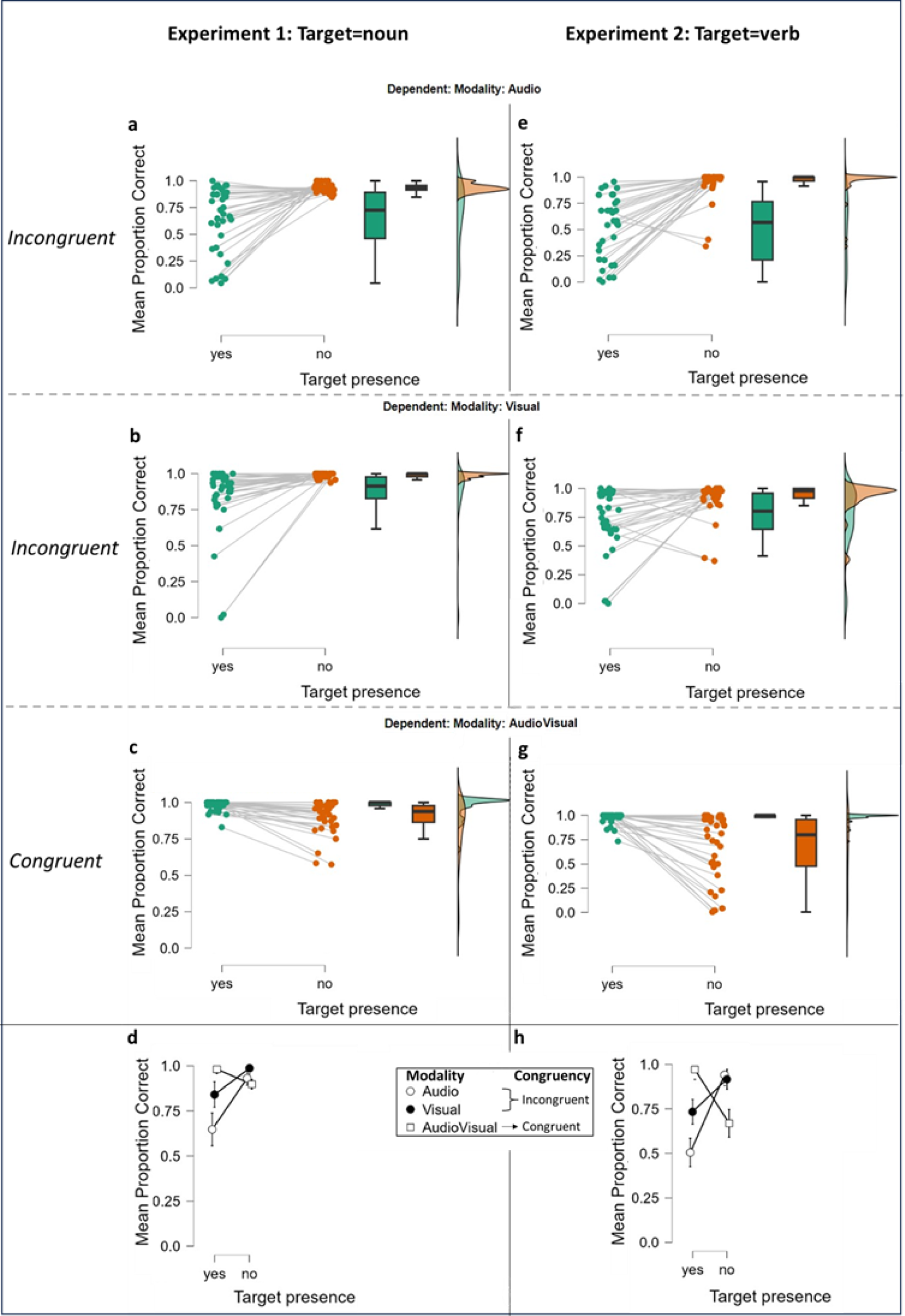
Mean proportion correct responses for target presence and target absence in Experiment 1 and Experiment 2. Box-and-whisker plots in (a), (b) and (c) represent the interquartile range (IQR) for Experiment 1 and in (e), (f) and (g) the IQR for Experiment 2; central, bold horizontal lines in (a), (b), (c) and (e), (f), (g) represent the medians; white squares, black circles and white circles in (d) and (h) represent the group mean proportion correct (Experiment 1: N=36, Experiment 2: N=34) when target (Target presence=yes) or target-related information (Target presence=no) was presented in the three modality conditions (audio, Visual, AudioVisual). Green and orange points represent the mean proportion of correct responses of individual participants. a) and e) show mean proportion correct individual scores in target present and target absent incongruent trials when target/target-related information was only heard for Experiment 1 and in Experiment 2, respectively. b) and f) show mean proportion correct individual scores in target present and target absent incongruent trials when target/target-related information was only seen for Experiment 1 and for Experiment 2, respectively. c) and g) show mean proportion correct individual scores in target present and target absent congruent trials (target/target-related information was both heard and seen) for Experiment 1 and for Experiment 2, respectively. d) and h) show mean proportion correct total scores in target present and target absent incongruent trials for Experiment 1 and for Experiment 2, respectively (target/target-related information was either only heard or only seen) vs. congruent trials (target/target-related information was both heard and seen).

*Figure 3c* shows results from movieclips with congruent information and the target noun being present in both modalities. Here, we observed the opposite pattern of results. When the target was present in both modalities, mean proportion of correct answers was high (M_audiovisual_= 0.981, SD_audiovisual_= 0.036) but was reduced in target absent trials (M_audiovisual_= 0.898, SD_audiovisual_= 0.111).

Note that in target absent trials, the target was completely absent in both modalities while target-related information was presented in one of the two modalities (incongruent movieclips) or in both modalities (congruent movieclips). Our results show that the presence of target-related information does not compromise participants’ performance in incongruent movieclips *(Figures 3a, 3b)*.

In a two-way repeated-measures ANOVA we report statistically significant main effects of Modality (F(2,70)=25.837, p< .001) and Target presence (F(1,35)=26.940, p< .001). The interaction between Modality and Target presence was also statistically significant (F(2,70)=20.265, p< .001). The post-hoc pairwise comparisons showed that during incongruent movieclips the performance was statistically worse when target or target-related information was presented in audio modality compared to visual (t=-5.567, p< .001) or audiovisual (t=-6.722, p<.001). Further post-hoc pairwise comparisons (see *Table 4a*) showed that the performance was superior during congruent movieclips (target present in both modalities) compared to incongruent movieclips (target present only through audio or only through vision) (t_Audio vs. AudioVisual_ =-9.094, p< .001; t_Visual vs. AudioVisual_ =-3.819, p= 0.002). We also observed that in incongruent movieclips the presence of target affects the accuracy in responses on a statistically significant level (t_AusdioYes vs. AudioNo_ =-7.044, p< .001; and for visual: t_VisualYes vs. VisualNo_ =-3.619, p= 0.004). We also found a correlation between participants’s age and performance (proportion of correct answers) -see *Figure A2* in Appendix.

### Experiment 2: Target is always a verb

The results indicate a statistically significant difference in mean proportion correct responses depending on the presence of the target and the modality in which the target or target-related information was presented. Figure 3e-f shows results from movieclips with incongruent information where the target verb is presented only in the auditory or visual modality. Similar to Experiment 1, we observed again a similar pattern of results: reduced accuracy in responses when the target was present only in audio (M_audio_= 0.505, SD_audio_= 0.311) and only in visual modality (M_visual_=0.734, SD_visual_= 0.280), whereas superior performance when the target was absent altogether and reduced variance in responses (M_audio_= 0.940, SD_audio_= 0.153; M_visual_= 0.916, SD_visual_= 0.150).

*Figure 3g* shows results from movieclips with congruent information and target verb being presented in audio and visual modality. We observed again the opposite pattern of results. When the target was present in both modalities mean proportion of correct answers was high (M_audiovisual_= 0.971, SD_audiovisual_= 0.059), whereas in target absent trials, the mean proportion of correct responses was reduced (M_audiovisual_= 0.668, SD_audiovisual_= 0.346).

Like in Experiment 1, results show that target-related information does not compromise participants’ performance in incongruent movieclips *(Figures 3e,3f)*.

In a two-way repeated-measures ANOVA we report statistically significant main effects of Modality (F(2,66)=19.728, p< .001) and Target presence (F(1,33)=12.812, p= 0.001). The interaction between Modality and Target presence was also statistically significant (F(2,66)=39.960, p< .001). The post-hoc pairwise comparisons showed that the performance was statistically worse when target or target-related information was presented in audio modality compared to visual (t=-5.576, p< .001) or audiovisual (t=- 5.293, p<.001). Regarding the comparison between congruent and incongruent conditions during target present trials, superior performance was noted when target was presented in both modalities compared to when it was present only through audio (t_Audio vs. AudioVisual_ =-10.180, p< .001) or only through vision (t_Visual vs. AudioVisual_ =-5.180, p< .001). However, during target absent trials, significantly lower performance was noted when the target-related information was presented in both modalities compared to unimodally (t_Audio vs. AudioVisual_ =5.940, p< .001; t_Visual vs. AudioVisual_ =5.408, p< .001). We also observed that in both congruent and incongruent movieclips the presence of target, affects the accuracy in responses on a statistically significant level (for audio: t_AusdioYes vs. AudioNo_ =-7.689, p< .001; for visual: t_VisualYes vs. VisualNo_ =-3.219, p= 0.009; for audiovisual: t_AusdiovisualYes vs. AudiovisualNo_=5.335, p< .001). We found no correlation between participants’s age and performance (proportion of correct answers) -see *Figure A2* in Appendix.

### Semantics in Mean Proportion Correct

### Experiment 1: Target is always a noun

We tested for the effect of semantics only in target absent trials, since only those included target-related information (verb or noun) either in one of the modalities (incongruent movieclips) or in both (congruent movieclips). As mentioned in previous sections, in target absent trials incongruent movieclips did not contain the target in either modality, but rather one modality included information (a verb or a noun) which was semantically related to the target. For example, the target “knife” was not presented in the visual clip nor heard in the audio sentence, rather the audio sentence included the verb “cut” (which is semantically related to the word “knife”). Respectively, in congruent movieclips the information presented in both modalities was semantically related to the target “knife”, i.e., participants were hearing the sentence “Someone cut the lemons” while watching two hands mixing a glass of juice with many lemon slices in the front ground.

We observed a different pattern when the target-related audio sentence included a semantically related noun (see Figure 4a-c). While the pattern of results is similar for the two congruent semantic conditions (M_noun_= 0.903, SD_noun_= 0.130 vs. M_verb_= 0.892, SD_verb_= 0.115), we observe superior performance for incongruent trials when the target-related information was available in the form of a noun (M_noun_= 0.994, SD_noun_= 0.015 vs. M_verb_= 0.867, SD_verb_= 0.075). A two-way repeated-measures ANOVA was conducted to test for the effects of congruency (two levels: incongruent/congruent) and semantics (two levels: verb/noun) on the accuracy of responses. In this case, testing for congruency effects means to test whether the semantically related audio sentences presented with their identical video clips (congruent movieclips) showed statistically significant difference in performance compared to when the semantically related audio sentences were presented together with a random video clip (incongruent movieclips). In our analysis, Congruency did not yield a significant result (F(1,35)=2.580, p=0.117). On the other hand, the main effect of Semantics (F(1,35)=29.877, p< .001) was found to be statistically significant, so did the interaction of Congruency and Semantics (F(1,35)=49.683, p< .001).

**Figure 4.**
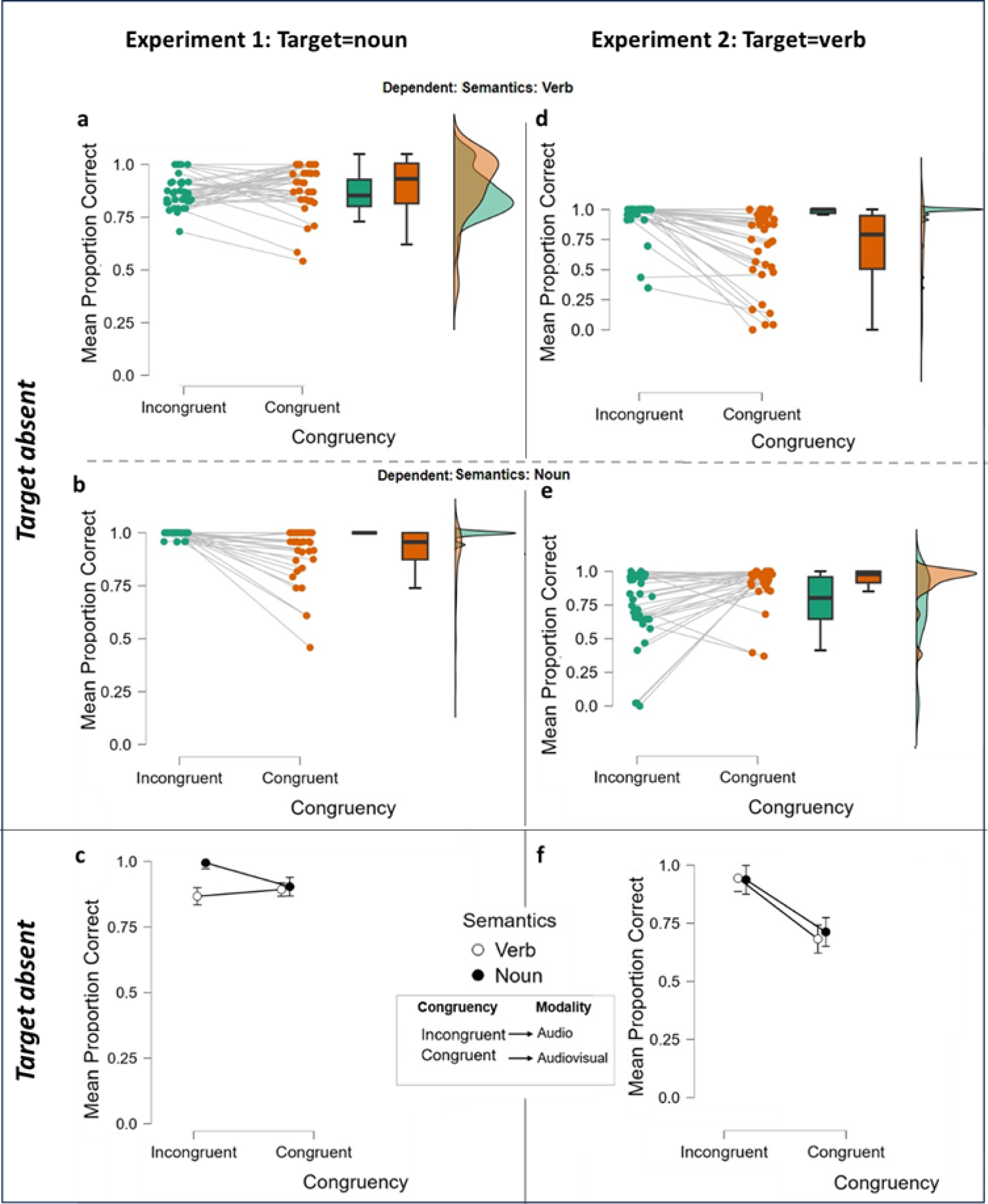
Mean proportion correct for incongruent and congruent semantic conditions in target absent trials in Experiment 1 and Experiment 2. Box-and-whisker plots in (a), (b) represent the interquartile range (IQR) for Experiment 1 and in (d), (e) the IQR for Experiment 2; central, bold horizontal lines in (a), (b) and (d), (e), represent the medians; black circles and white circles in (c) and (f) represent the group mean proportion correct (Experiment 1: N=36, Experiment 2: N=34). Green and orange points represent the mean proportion of correct responses of individual participants. a) and d) show mean proportion correct individual scores in target absent trials when target-related audio sentences included a verb accompanied by a semantically incongruent visual clip vs. a semantically congruent visual clip for Experiment 1 and in Experiment 2, respectively. b) and e) show mean proportion correct individual scores in target absent trials when target-related audio sentences included a noun accompanied by a semantically incongruent visual clip vs. a semantically congruent visual clip for Experiment 1 and in Experiment 2, respectively. c) and f) show mean proportion correct total scores in incongruent trials for verb vs. noun (audio sentence with a target related verb vs. noun accompanied by a semantically incongruent visual clip) vs. congruent trials for Experiment 1 and in Experiment 2, respectively (audio sentence with a target related verb vs. noun accompanied by a semantically congruent visual clip).

Post-hoc tests revealed that the observed superior performance for incongruent trials when the target-related sentence included a noun compared to when it included a verb was statistically significant (t_verb vs. noun_=-8.428, p< .001). Moreover, semantically target-related noun was associated with superior accuracy in responses when presented during incongruent movieclips compared to when it was presented during congruent movieclips (t_InNoun vs. CoNoun_ =4.144, p< .001). Finally, a statistically significant difference was noted between target-related verb when presented during congruent movieclips and target-related noun when presented during incongruent movieclips (t_CoVerb vs. InNoun_=-4.256, p< .001).

### Experiment 2: Target is always a verb

Similar to Experiment 1, we tested for the effect of semantics only in target absent trials in incongruent movieclips, which did not contain the target in either modality, but only a semantically related word (verb or noun). For example, the target “drink” was not presented in the visual clip nor heard in the audio sentence, rather the audio sentence included the noun “juice” (which is semantically related to the word “drink”). Respectively, in congruent movieclips the information presented in both modalities was semantically related to the target “drink”, i.e., participants were hearing the sentence “She is drinking a lemonade” while watching a pregnant woman drinking a lemonade in a tavern while a man was sitting beside her.

As shown in Figure 4d-f, we observe similar patterns of results when the target-related audio sentence included a semantically related noun or verb in incongruent and congruent movieclips (incongruent: M_noun_= 0.937, SD_noun_= 0.156 vs. M_verb_= 0.944, SD_verb_= 0.152; congruent: M_noun_= 0.712, SD_noun_= 0.335 vs. M_verb_= 0.682, SD_verb_= 0.323). A two-way repeated-measures ANOVA was conducted to test for the effects of congruency (incongruent/congruent) and semantics (verb/noun) on the accuracy of responses. In contrast to Experiment 1, the main effect of congruency was found to be statistically significant (F(1,33)=23.292, p< .001), as well as the interaction of Congruency and Semantics (F(1,33)=7.219, p= 0.011), whereas semantics did not yield a significant result (F(1,33)=3.132, p= 0.086). Post-hoc tests revealed that performance in incongruent movieclips was almost impeccable compared to congruent movieclips, regardless of the presentation of a semantically target-related verb or noun (t_InVerb vs. CoVerb_=5.146, p< .001; t_InNoun vs. CoNoun_=4.417, p< .001). Generally, when the target-related verb was presented audiovisually performance was at its worst (t_InVerb vs. CoNoun_=4.554, p< .001; t_CoVerb vs. InNoun_=- 5.016, p< .001; t_CoVerb vs. CoNoun_=-3.1644, p= 0.005), showing that when participants tried to detect the target verb, the crossmodal presentation of a target-related verb confused their decision.

### Reaction Times

### Experiment 1: Target is always a noun

We observed the fastest mean responses in congruent trials when the target was present audiovisually (M_audiovisual_= 0.520s, SD_audiovisual_=0.230). The second fastest scores were observed for the trials where the target was present only in the audio modality (M_audio_=0.590s, SD_audio_=0.257), whereas slower responses were shown in trials where the target was presented only in the visual modality (M_visual_=0.620s, SD_visual_=0.358). In target absent trials, mean RTs were generally slower compared to when the target was present. Here, faster responses were recorded when the target-related stimulus appeared in the visual modality (M=0.670s, SD=0.302), followed by slower responses for the audio modality (M=0.683s, SD=0.317) and the slowest when presented in both modalities (M=0.741s, SD=0.366). The two-way repeated measures ANOVA showed that the performance regarding mean RTs was not associated with statistically significant differences between modality conditions (F(2,70)=0.384, p=0.683). However, there was a statistically significant main effect of Target presence (F(1,35)=20.825, p<.001) and a statistically significant interaction between Modality and Target presence (F(2,70)=8.461, p< .001). The post-hoc pairwise comparisons showed that the performance was statistically worse when target-related information was presented in both modalities (see *Table 5a*). Finally, we found a correlation between participants’s age and reaction times -see *Figure A2* in Appendix.

**Table 5a.** Post Hoc Comparisons for Mean RT in target present and target absent conditions (Modality * Target presence) in Experiment 1.

|  |  | Experiment 1 |  |  |  |
| --- | --- | --- | --- | --- | --- |
| | | Mean Difference | SE | t | $P_{\text{holm}}$ |
| Audio, yes | Visual, yes | -0.031 | 0.027 | -1.120 | 0.529 |
|  | Audiovisual, yes | 0.069 | 0.027 | 2.543 | 0.097 |
|  | Audio, no | -0.093 | 0.036 | -2.554 | 0.097 |
|  | Audiovisual, no | -0.152 | 0.034 | -4.494 | < .001 |
| Visual, yes | Audiovisual, yes | 0.100 | 0.027 | 3.663 | 0.004 |
|  | Visual, no | -0.050 | 0.036 | -1.375 | 0.518 |
|  | Audiovisual, no | -0.121 | 0.034 | -3.589 | 0.006 |
| Audiovisual, yes | Audio, no | -0.162 | 0.034 | -4.812 | < .001 |
|  | Visual, no | -0.150 | 0.034 | -4.443 | < .001 |
|  | Audiovisual, no | -0.221 | 0.036 | -6.059 | < .001 |
| Audio, no | Visual, no | 0.012 | 0.027 | 0.458 | 0.648 |
|  | Audiovisual, no | -0.059 | 0.027 | -2.148 | 0.168 |
| Visual, no | Audiovisual, no | -0.071 | 0.027 | -2.606 | 0.092 |

**Figure 5.**
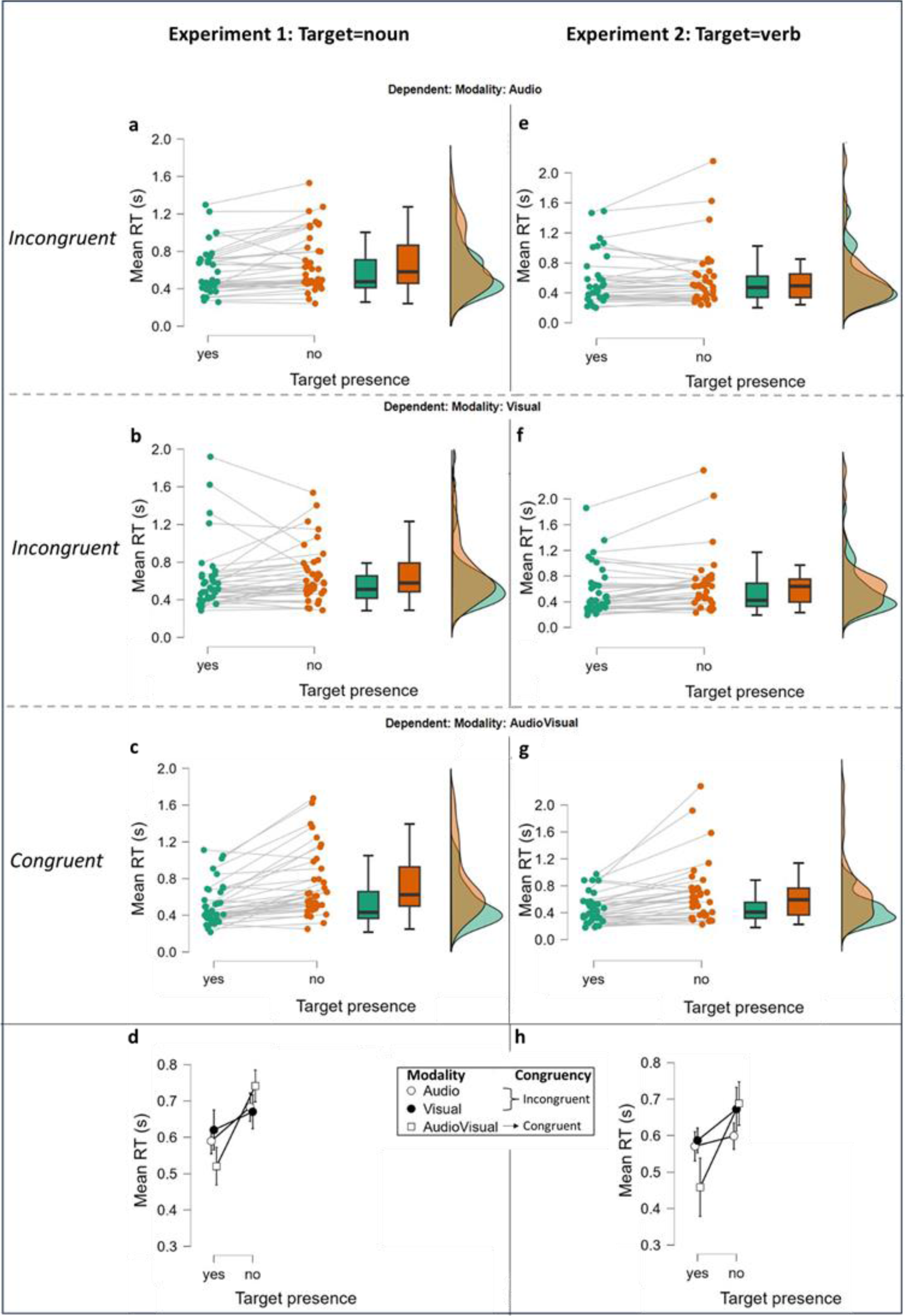
Mean RT scores for target presence and target absence in Experiment 1 and Experiment 2. Box-and-whisker plots in (a), (b) and (c) represent the interquartile range (IQR) for Experiment 1 and in (e), (f) and (g) the IQR for Experiment 2; central, bold horizontal lines in (a), (b), (c) and (e), (f), (g) represent the medians; white squares, black circles and white circles in (d) and (h) represent the group mean RT scores (Experiment 1: N=36, Experiment 2: N=34) when target (Target presence =yes) or target-related information (Target presence = no) was presented in the three modality conditions (audio, Visual, AudioVisual). Green and orange points represent the mean RT scores of individual participants. Note that range in (d) and (h) is 0.3 to 0.8. a) and e) show mean RT individual scores in target present and target absent incongruent trials when target/target-related information was only heard for Experiment 1 and in Experiment 2, respectively. b) and f) show mean RT individual scores in target present and target absent incongruent trials when target/target-related information was only seen for Experiment 1 and for Experiment 2, respectively. c) and g) show mean RT individual scores in target present and target absent congruent trials (target/target-related information was both heard and seen) for Experiment 1 and for Experiment 2, respectively. d) and h) show mean RT total scores in target present and target absent incongruent trials for Experiment 1 and for Experiment 2, respectively (target/target-related information was either only heard or only seen) vs. congruent trials (target/target-related information was both heard and seen).

### Experiment 2: Target is always a verb

We observed the fastest mean responses when the target was present audiovisually (M_audiovisual_= 0.459s, SD_audiovisual_=0.216). The mean RT scores when the target was present either only in audio or visual modality were almost similar (M_audio_=0.571s, SD_audio_=0.340; M_visual_=0.587s, SD_visual_=0.376). As in Experiment 1, in target absent trials the mean RTs were generally slower compared to when the target was present. This time, faster responses were recorded when the target-related stimulus appeared in the audio modality (M=0.599s, SD=0.405), whereas slower responses were observed when the target-related stimulus was presented in the visual modality (M=0.672s, SD=0.467) and the slowest again when presented in both modalities (M=0.688s, SD=0.462). The two-way repeated measures ANOVA showed that the performance regarding mean RTs was associated with statistically significant differences between modality conditions (F(2,66)=3.533 p=0.035), as well as target presence conditions (yes/no) (F(1,33)=11.117, p= 0.002), and the interaction between Modality and Target presence (F(2,66)=13.595, p< .001) was also statistically significant. Post Hoc pairwise comparisons (see *Table 5b*) showed that performance was faster when the target was presented in both modalities (congruent movieclips) compared to when it was presented in incongruent movieclips (t_Audio vs. AudioVisual_ =3.747, p= 0.003; t_Visual vs. AudioVisual_ =4.280, p< .001). They also revealed slowest responses in congruent movieclips when the target was absent compared to when it was present (t =-5.557, p< .001). Finally, similar to Experiment 1, we found a correlation between participants’s age and reaction times.

**Table 5b.**
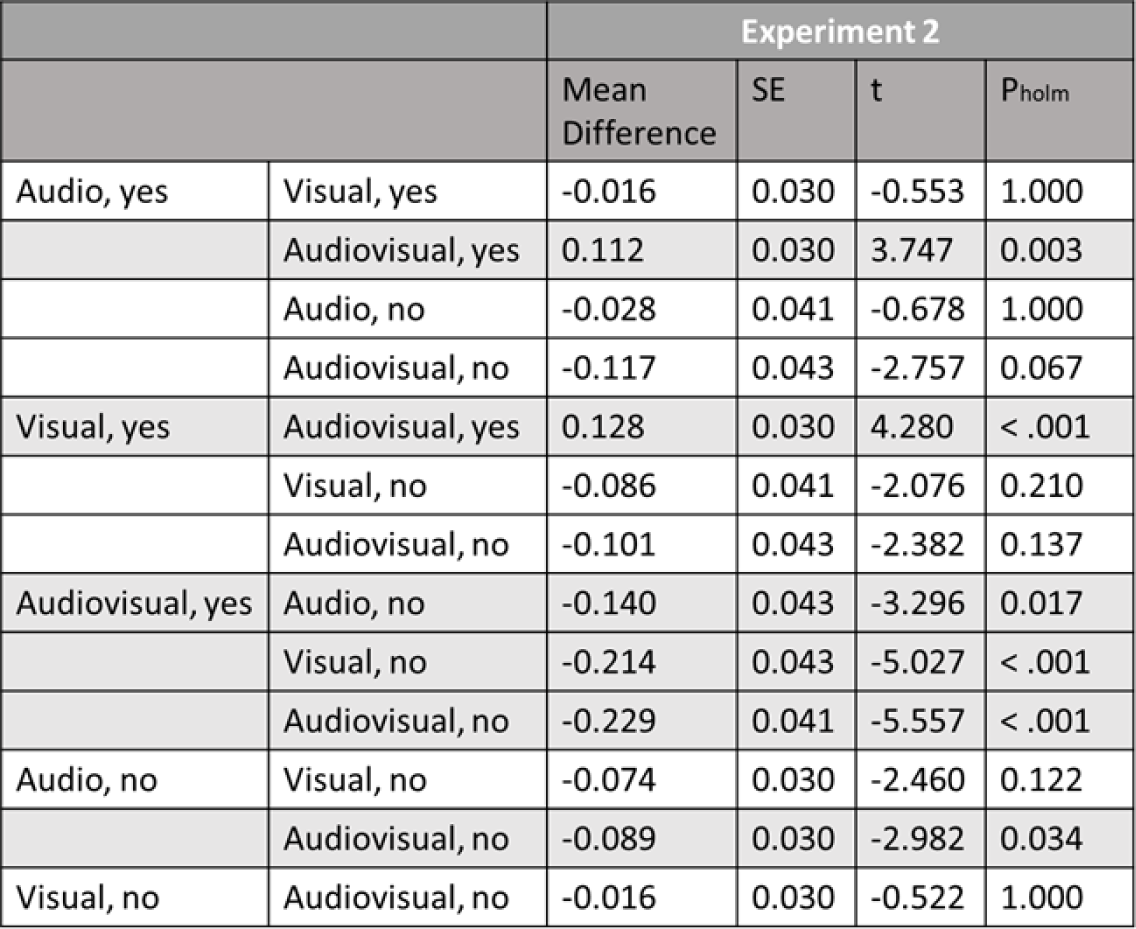
Post Hoc Comparisons for Mean RT in target present and target absent conditions (Modality * Target presence)

### Semantics in RTs

### Experiment 1: Target is always a noun

The fastest mean RT scores were found in incongruent movieclips regardless of whether they included a semantically related noun (M_noun_=0.684s, SD_noun_=0.336) or a verb (M_verb_=0.681s, SD_verb_=0.339), compared to congruent movieclips (M_verb_=0.726s, SD_verb_=0.352; M_noun_=0.758s, SD_noun_=0.386).

A two-way repeated-measures ANOVA was conducted in target absent trials to test for the effects of congruency and semantics on mean RTs. A significant effect was found for Congruency (F(1,35)=5.908, p=0.020), but not for Semantics (F(1,35)=0.782, p=0.382) or the interaction between the two factors (F(1,35)=0.410, p=0.526).

**Figure 6.**
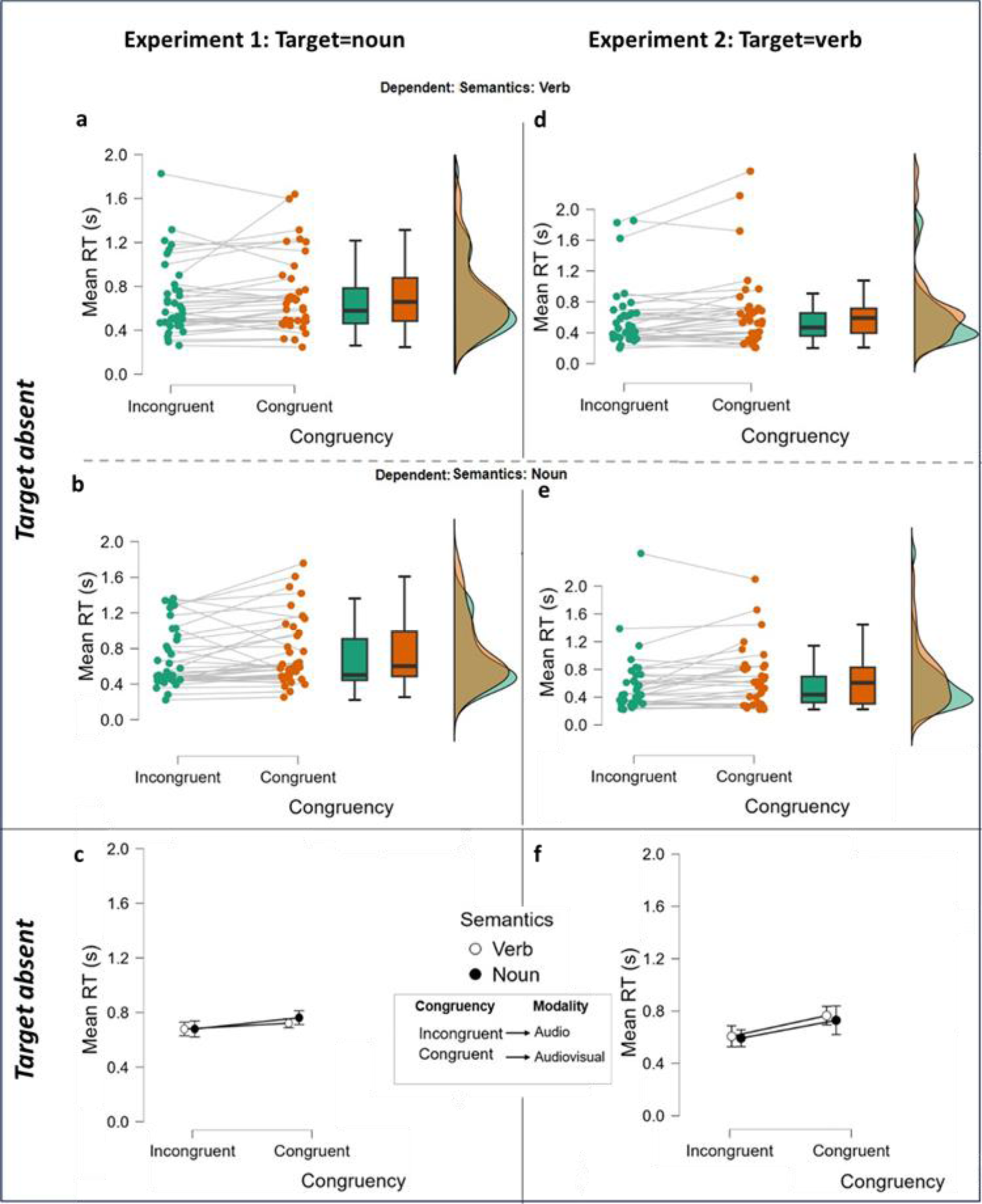
Mean RT scores for incongruent and congruent semantic conditions in target absent trials in Experiment 1 and Experiment 2. Box-and-whisker plots in (a), (b) represent the interquartile range (IQR) for Experiment 1 and in (d), (e) the IQR for Experiment 2; central, bold horizontal lines in (a), (b) and (d), (e), represent the medians; black circles and white circles in (c) and (f) represent the group mean RT (Experiment 1: N=36, Experiment 2: N=34). Green and orange points represent the mean RT of individual participants. a) and d) show mean RT individual scores in target absent trials when target-related audio sentences included a verb accompanied by a semantically incongruent visual clip vs. a semantically congruent visual clip for Experiment 1 and in Experiment 2, respectively. b) and e) show mean RT individual scores in target absent trials when target-related audio sentences included a noun accompanied by a semantically incongruent visual clip vs. a semantically congruent visual clip for Experiment 1 and in Experiment 2, respectively. c) and f) show mean RT total scores in incongruent trials for verb vs. noun (audio sentence with a target related verb vs. noun accompanied by a semantically incongruent visual clip) vs. congruent trials for Experiment 1 and in Experiment 2, respectively (audio sentence with a target related verb vs. noun accompanied by a semantically congruent visual clip).

### Experiment 2: Target is always a verb

Similar to Experiment 1, the fastest mean RT scores were found in incongruent movieclips (M_noun_=0.586s, SD_noun_=0.429; M_verb_=0.604s, SD_verb_=0.410), compared to congruent movieclips (M_noun_=0.673s, SD_noun_=0.435; M_verb_=0.699s, SD_verb_=0.510).

The two-way repeated-measures ANOVA revealed a significant effect for Congruency (F(1,33)=11.723, p=0.002), but not for Semantics (F(1,33)=0.789, p=0.381) or the interaction between the two factors (F(1,33)=0.029, p=0.867).

## Discussion

The present study was conducted to examine audiovisual integration in the perception of synchronous, and semantically congruent and incongruent audiovisual stimuli that include movement and spoken sentences. Specifically, we investigated the role of semantic associations in the perception of congruent and incongruent crossmodal stimuli. Accuracy scores indicated a statistically significant difference between modalities in which the target or target-related information was presented, as well as between target present conditions. In accordance with previous studies (Barrett & Newell, 2015; Smayda et al., 2016; Brooks et al., 2018), we also report a significant negative correlation between participants’ age and accuracy scores for both experiments, as well as between participants’ age and mean RTs only for Experiment 1 (where target is always a noun). We provide for the first time evidence showing that in congruent target absent trials where the information is presented audiovisually, accuracy is significantly compromised when target related information is a verb.

### Target Present Trials

In our study, judgments for targets’ presence revealed that listening to spoken sentences that include the target stimuli while perceiving incongruent visual information was associated with lowest accuracy in responses. In an earlier study, Soto-Faraco et al. (2002) studied the dynamic capture of audiovisual motion information using sounds and led lights that either had the same or opposite direction, and suggested the existence of an audiovisual dynamic capture effect in order to explain the modulation of the perceived direction of auditory apparent motion caused by apparent motion in visual modality. This alternation of audio information has been reported in many studies where simple non-semantic audiovisual stimuli were presented with spatial incongruency between the auditory and visual signals, such as this of the ventriloquism effect (Bruns, 2019) as well as the McGurk effect, but has been also reported in studies that used semantic incongruent audiovisual stimuli (Opoku-Baah et al., 2021). Thus, our finding is consistent with previous studies that examined the visual influences on auditory perception and adds another aspect, this of complexity of information content, in perceptual alterations of audio semantic information due to their simultaneous presentation with conflicting but task-relevant semantic visual information. Our study revealed also that watching a video scene, where the target stimuli appeared while listening to incongruent spoken sentences, was also associated with lower accuracy in judgments. These findings provide support for the suppressive effect that semantic sound can have on visual perception (Chen & Spence, 2010; Keetels & Vroomen, 2011; Bolognini et al., 2013; Fujisaki et al., 2014; Ortega et al., 2014; Hidaka & Ide, 2015; Hsiao et al., 2012; Williams et al., 2022). Regarding this effect, Hidaka and Ide (2015) showed decreased performance during visual orientation discrimination tasks due to the appearance of spatially and temporally congruent sounds (white noise bursts) through headphones. They explained their findings on the basis of potentially direct and close interactions of neural responses occurring across modalities during the observation of audiovisual stimuli.

Moreover, the results in both experiments indicate the complementary role of congruent audiovisual information in perceiving target stimuli when presented in a crossmodal interface. In target present trials, performance was significantly higher with semantically congruent audiovisual stimuli (compared with incongruent audiovisual stimuli) and this has been previously reported (Laurienti et al., 2004; Molholm et al, 2004; Chen & Spence, 2010; Xie et al., 2017; Rekow et al., 2022). It is somewhat surprising that reaction times did not differ significantly between audiovisual and audio modality in Experiment 1. Molholm et al. (2004), for example, observed a trend of significantly higher performance in an object detection task, where a combined influence of crossmodal inputs (i.e. line-drawing pictures and vocalizations of animals) was suggested. They reported not only significantly higher accuracy but also significantly faster target identification in congruent conditions when the picture and vocalization of the same animal were matched, compared to when the target was presented only in one sensory modality. To date, it is strongly supported that this significant difference in RTs is due to the semantic congruency between the two modality signals (Laurienti et al., 2004; Molholm et al, 2004; Mastroberardino, Santangelo, & Macaluso, 2015; Tsilionis & Vatakis, 2016). However, Letts, Basharat, and Barnett-Cowan (2022) bring the parameter of valence being another significant factor in multisensory integration. Our results provide further evidence that semantic congruency cannot be a critical factor on its own for determining multisensory perception. We propose that the complexity of language structure (i.e., word or sentence) and its relation to the target stimulus may play the most important role, especially in real-word multimodal perception.

### Target Absent Trials

In all trials where the target was absent, we included for the first time target-related information instead of completely irrelevant information as previous studies did. Although these trials included target-related information, this information did not compromise participants’ performance. Specifically in both experiments, the modality in which the target-related information appeared did not influence the performance significantly, as performance was near veridical whether target-related information was presented through vision or audition alone. As discussed so far, incongruent audio and visual information interact with each other to create coherent representation of that information (Roach, Heron, & McGraw, 2006; Tsilionis & Vatakis, 2016). Thus, during incongruent crossmodal signals, even when the presented information in either modality is related to the target object, word or concept, the coherent representation induced by the crossmodal perceptual process remains accurate to detect the absence of the target. However, it is possible that during these trials the suppression effect is still affecting the performance, but this time by positively interfering with task demands, meaning that the suppression effect may overlap the target-related information.

On the other hand, in the case of congruent movieclips, we observe significant decrease in mean RTs in both experiments, and decrease in judgments in Experiment 1 (when the target was a noun), which was significant in Experiment 2 (when the target was a verb). One possibility is that participants’ judgements may have been affected by the relation between congruent crossmodal information and target-related information which in our analysis was found to be statistically significant. The presentation of target-related information in both modalities may result in a confusion or even an illusion of what has been seen and heard. This confusion may arise from the combination of two factors: 1) strong audiovisual integration that is formed temporally for each congruent trial, and its strength is due to the semantic congruency of audiovisual information, together with 2) strong semantic schemata/concepts that exist between objects, locations, actions, movements etc. As discussed so far regarding the first factor (1), congruent audiovisual stimuli tend to build stronger connections between the perceived information and therefore facilitate perception. These strong connections have been also indicated through their efficient neural representations in the neuroanatomical surface (Li et al., 2011). As for the second factor (2): The strength of semantic concepts has been well-established for settings that examine attention performance such as visual search tasks, where research indicates the significant influence of semantic concepts even when there is low accuracy of the concept detectors (Long & Chang, 2014). In addition to this, a strong context-dependent association of audiovisual integration with multiple interactions in various brain regions has been reported (Diaconescu, Alain, & McIntosh, 2011; Gao et al, 2022). In this matter, by combining factors (1) and (2) we could assume that semantics could create an effect of semantic relatedness on perceptual performance in congruent audiovisual movieclips depending on features such as the complexity of combined semantic information, the expectations of the observer, etc. which in turn activates more complex integrated brain processes. The slower response times in our results when target-related information was presented in congruent movieclips compared to when the target was, could be another evidence of this effect.

### The Role of Semantics (Verb vs. Noun) in Target Absent Trials

Furthermore, our experiment examined whether hearing a sentence that includes a word which is semantically related to the target stimuli, could affect judgements in incongruent and congruent movieclips. To date, it is the first study that used semantically related ‘distractors’ and replaced target absent trials with target-related (‘distractor’) trials. We grouped participant responses based on the type of the semantically target-related word that the audio sentences included: verbs or nouns. The results for both experiments indicate that in congruent movieclips, regardless of whether they included a semantically target-related verb or noun, participant performance was similar, with the worst judgment scores observed when the target was a verb (Experiment 2) and was accompanied with slower responses in both experiments. These results suggest that the potential confusion during congruent movieclips in target absent trials, which was described above to explain the decrease in performance, was not associated with the type of semantics of the presented stimuli (whether a semantically target-related verb or noun was presented). On the contrary the potential confusion might be associated with the type of the target (whether it was a verb or a noun), and the worse judgment scores in Experiment 2 (where target was always a verb) could indicate the higher difficulty level of this task compared to this of Experiment 1 where the target was always a noun.

Moreover, when the target was a noun (Experiment 1) in incongruent movieclips while listening to a sentence that includes a semantically related noun performance was almost impeccable (M=99.4%), compared to listening to a sentence that includes a semantically related verb. The first condition seems to agree with the general results of incongruent movieclips in target absent trials. Respective to what has been previously mentioned, this pattern could suggest that target-related nouns did not work as distractors for correctly judging the absence of the target in incongruent movieclips. This might be due to the fact that participants were able to make easier comparisons between the noun that they have heard while seeing an irrelevant visual clip and the target noun that they were looking to hear and/or see. Interestingly, this was not the case in Experiment 2 when target-related verbs were presented. In general, nouns differ from verbs in the information level they can add in a sentence, but both cooperate to establish neural representations of objects and events (Faroqi-Shah, Sebastian, & Woude, 2018). For instance, nouns are related to objects, or subjects who perform actions, and can complete the meaning of actions, whereas verbs refer to actions and events, including also -in many languages-temporal information about the actions, and thus indicating the syntactic complexity of verbs (Geng et al., 2022; de Aguiar & Rofes, 2022). According to King and Gentner (2019), semantic context adaptations for verbs show to be driven by online adjustments whereas for nouns by sense-selection. Maguire et al. (2015) provide evidence for higher neural activity demands in action-verb based identification compared to object-noun. In addition to this, interesting assumptions arise by theories of semantic change regarding reinterpretation or form-meaning remapping of listeners depending on task demands (Dubossarsky, Weinshall, & Grossman, 2016). A frequent observation of verbs’ meaning adaptations has been reported which does not depend on the polysemy of verbs, rather on semantic strain contexts (King & Gentner, 2019). Taking these into consideration, we can propose that our results may at some level provide further evidence of the semantic complexity of verbs and the flexible nature of their cognitive representations compared to the simpler and more stable nature of nouns.

### General Comments and Limitations

The findings of the current study can be implemented in various settings. Education approaches already apply methods that not only take advantage of the enhancement effect induced by audiovisual information (Opoku-Baah et al., 2021), but also consider the semantic associations that words carry. For instance, when introducing a difficult topic or vocabulary to the students, educators use the Keyword Method in order to help students memorize the new word or content by visualizing it i.e. building visual representations (Chou, 2012; Dolean, 2014). We suggest that such educational approaches should give weight to the impact of semantic complexity, and the role of noun synonyms not only in helping memorizing but also in accurately perceiving complex information. Following, based on previous behavioural studies that report the influence of semantics in decision-making responses, our findings could in turn give rise to future research in AI design. Precisely, research in the fields of computational intelligence and AI is aiming at developing computational approaches that can be used for learning by perceiving information from audio and visual modalities in such a way as human perception works (Wei et al., 2023). The present findings add to the development of such computational approaches highlighting the influence of semantic relatedness and complexity between information provided from audio and visual modalities.

Furthermore, it is important to mention that our analysis did not examine the movement factor, which was included in the real-word based movieclips, since our design did not give weight to effects of movement on crossmodal-based behavioural responses. However, research focusing on language accounts, suggests that meanings of words depend on their perceptual and motor representations (Faroqi-Shah, Sebastian, & Woude, 2018) Thus, it may be worthy to further investigate for potential influence of movement in perceived audiovisual semantic information and test whether motor representations linked with words influence in any way audiovisual integration.

## Conclusion

Our study tested for existence of semantic congruency effects, as well as the appearance of a target either only in the visual clip (video) or only in the audio sentence or in both influences the performance of human participants in target detection judgments. These results were compared with those of target absent trials, which included target-related information either only in the visual clip/the audio sentence or in both. Our findings come in alignment with previous research that support enhancement in judgments when semantically congruent audiovisual information is simultaneously presented. Moreover, we provide further evidence of the effect of audio modality on visual and vice versa during crossmodal perceptual integration when using movieclips and, for the first time, audio sentences. To date it is also the first time a study involves target-related information to test how this may affect the perceptual integration process, in particular accuracy in judgments and RT. Thus, our approach focused on semantic congruency effects and complexity of semantics in target absent trials in order to examine whether seeing a target-related video accompanied by an audio sentence that includes a target-related verb vs. noun could impact performance. We tested this hypothesis also in comparison with the incongruent condition (non-target visual clip accompanied with audio sentence including target-related verb vs. noun). Our results indicate the critical role of complexity of semantics in crossmodal perceptual integration and could further support the assumption of cognitive representations induced by verbs being more elastic, and therefore, concluding in more complex associations with other verbs, nouns, objects, locations etc. compared to the simpler and more stable cognitive representations that depend on nouns. Our findings could enrich educational methods and multisensory learning techniques, and inspire computational and AI model design and learning.

## Data Availability Statement

Data for the main findings of this study are available at: https://osf.io/y3e4s/?view_only=1c4aeb77ef7843669155007bc0dcbcbe

## Appendix

**Table A1a.**
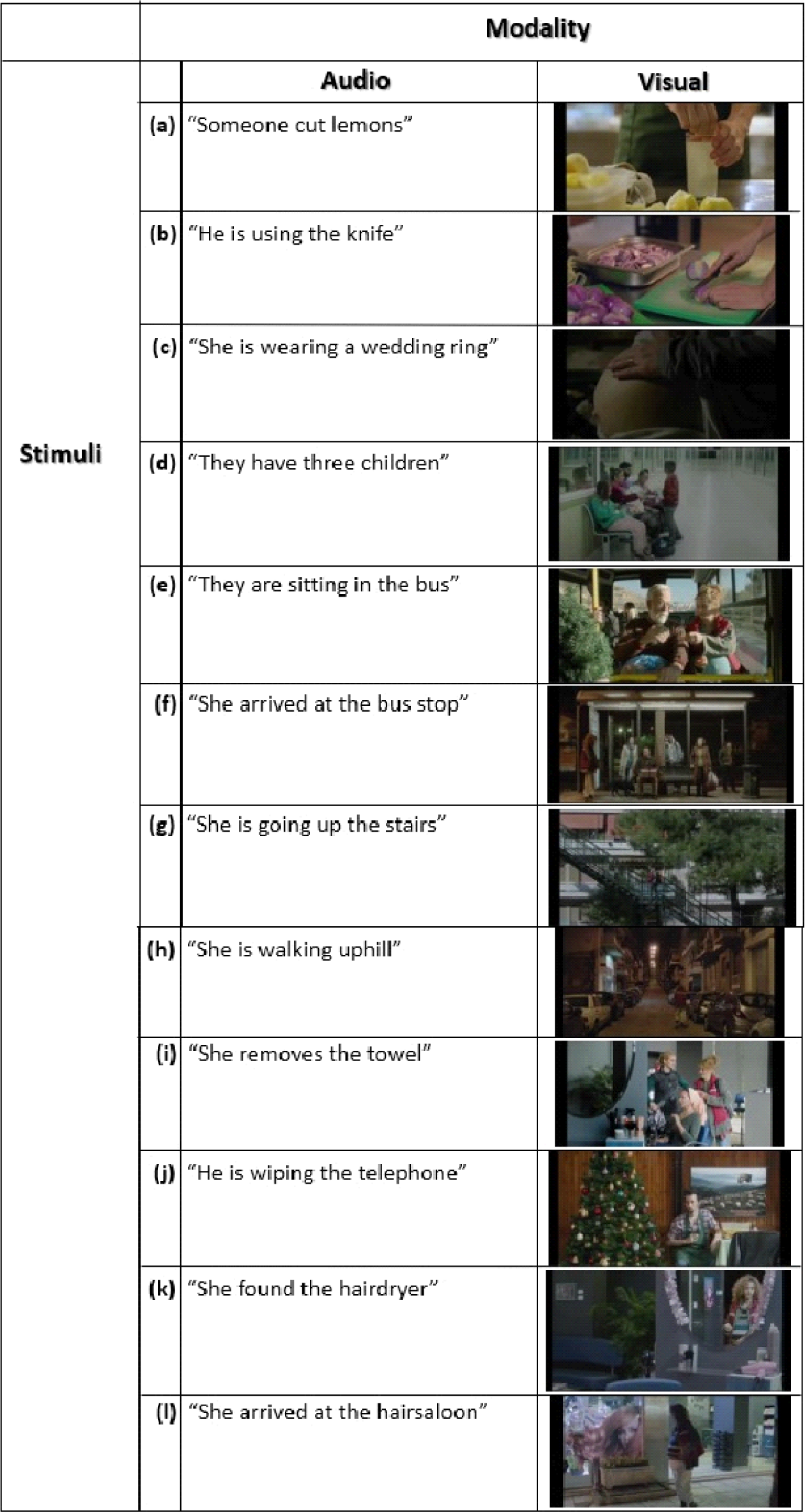
The audio and visual stimuli in Experiment 1.

**Table A2b.**
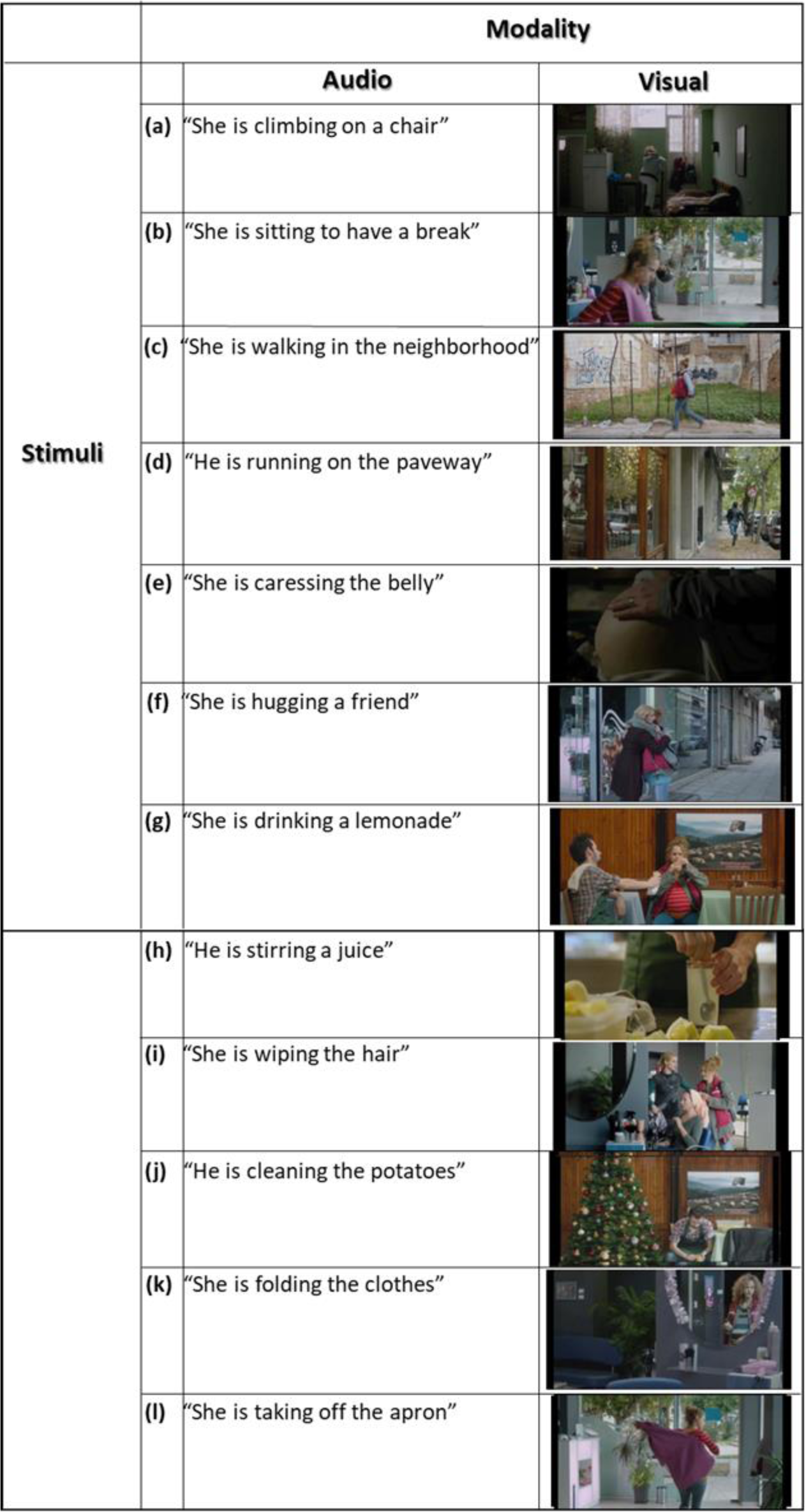
The audio and visual stimuli in Experiment 2.

Table A1b. The audio and visual stimuli in Experiment 2
|  | Modality |  |
| --- | --- | --- |
|  | Audio | Visual |
| **Stimuli** | (a) "She is climbing on a chair" |  |
|  | (b) "She is sitting to have a break" |  |
|  | (c) "She is walking in the neighborhood" |  |
|  | (d) "He is running on the paveway" |  |
|  | (e) "She is caressing the belly" |  |
|  | (f) "She is hugging a friend" |  |
|  | (g) "She is drinking a lemonade" |  |
|  | (h) "He is stirring a juice" |  |
|  | (i) "She is wiping the hair" |  |
|  | (j) "He is cleaning the potatoes" |  |
|  | (k) "She is folding the clothes" |  |
|  | (l) "She is taking off the apron" |  |

### Appendix Figures

**Figure A2.**
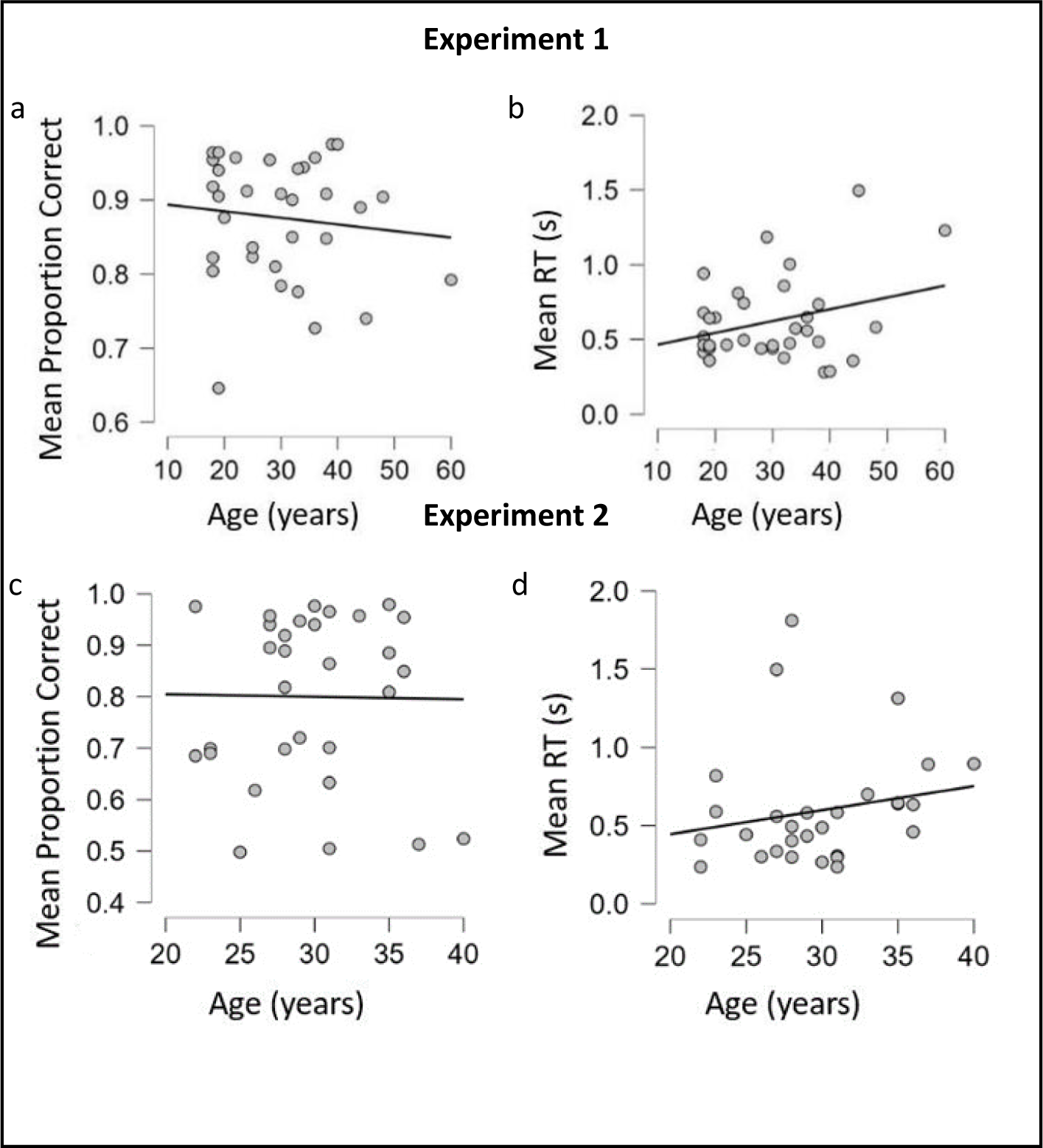
Correlation plots of mean proportion correct and mean RT vs. age, Experiment 1 and Experiment 2. Mean proportion correct responses and mean RTs for age, respectively. The lines in (a), (b) and (c), (d) represent the direction of means of total sample for Experiment 1 and Experiment 2 respectively, that is how much the y value (mean proportion correct/mean RT) increases or decreases across the x value (age). Grey points represent the means of individual participants. a) and c) show the correlation between total sample’s (*Experiment 1: N=36, Experiment 2: N=34*) mean proportion correct scores and age. b) and d) show the correlation between total sample’s mean RT scores and age.

**Figure A3.**
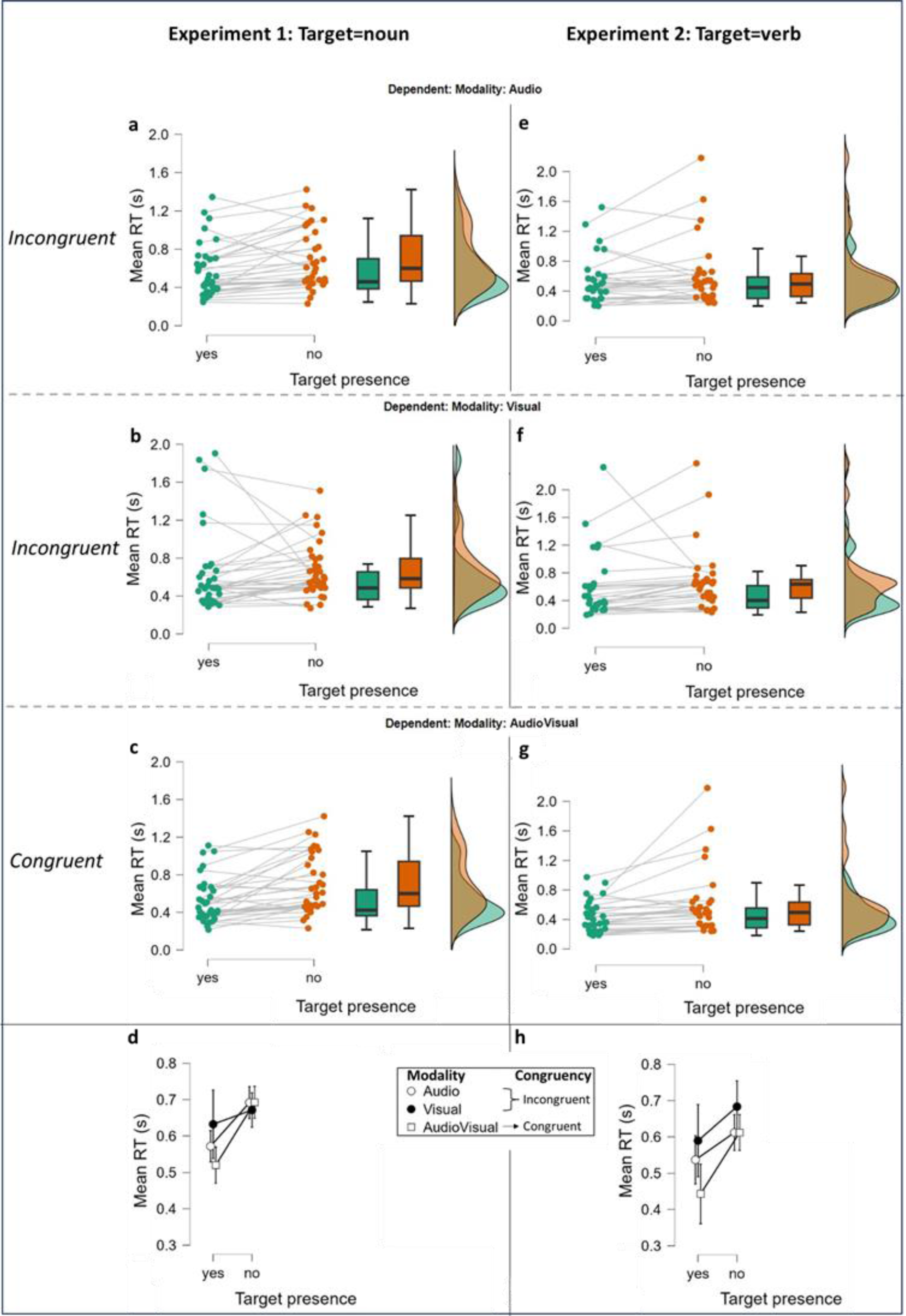
Mean RT scores for target presence and target absence in Experiment 1 and Experiment 2 (only correct answers). Mean RTs for target presence and target absence, analyzing only trials that were correctly answered. Box-and-whisker plots in (a), (b) and (c) represent the interquartile range (IQR) for Experiment 1 and in (e), (f) and (g) the IQR for Experiment 2; central, bold horizontal lines in (a), (b), (c) and (e), (f), (g) represent the medians; white squares, black circles and white circles in (d) and (h) represent the group mean RT scores (Experiment 1: N=36, Experiment 2: N=34) when target (Target presence=yes) or target-related information (Target presence=no) was presented in the three modality conditions (audio, Visual, AudioVisual). Green and orange points represent the mean RT scores of individual participants. Note that range in (d) and (h) is 0.3 to 0.8. a) and e) show mean RT individual scores in target present and target absent incongruent trials when target/target-related information was only heard for Experiment 1 and in Experiment 2, respectively. b) and f) show mean RT individual scores in target present and target absent incongruent trials when target/target-related information was only seen for Experiment 1 and for Experiment 2, respectively. c) and g) show mean RT individual scores in target present and target absent congruent trials (target/target-related information was both heard and seen) for Experiment 1 and for Experiment 2, respectively. d) and h) show mean RT total scores in target present and target absent incongruent trials for Experiment 1 and for Experiment 2, respectively (target/target-related information was either only heard or only seen) vs. congruent trials (target/target-related information was both heard and seen).

**Figure A4.**
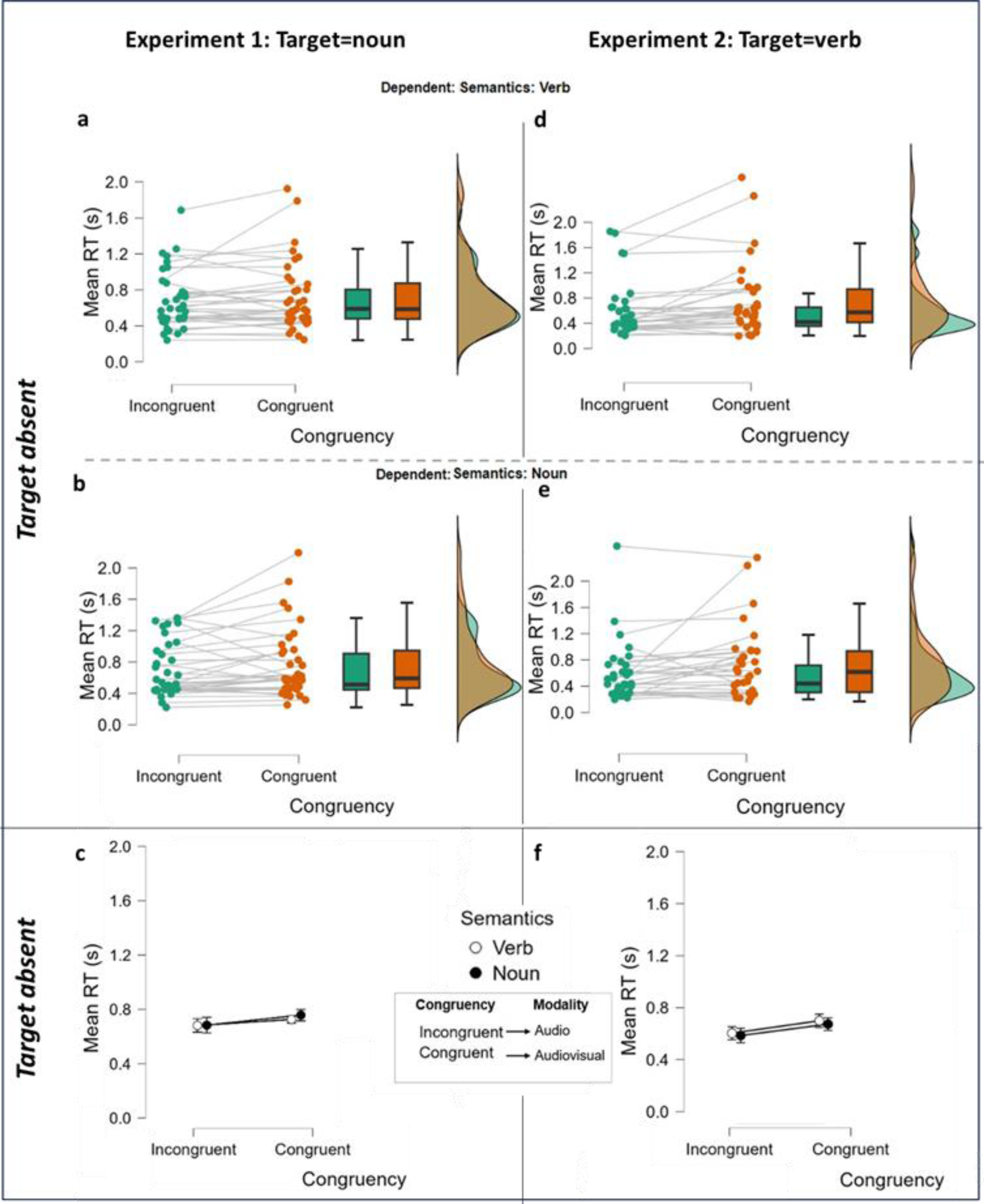
Mean RT scores for incongruent and congruent semantic conditions in target absent trials, Experiment 1 and Experiment 2 (only correct answers). Mean RTs for incongruent and congruent semantic conditions in target absent trials, analyzing only trials that were correctly answered. Box-and-whisker plots in (a), (b) represent the interquartile range (IQR) for Experiment 1 and in (d), (e) the IQR for Experiment 2; central, bold horizontal lines in (a), (b) and (d), (e), represent the medians; black circles and white circles in (c) and (f) represent the group mean RT (Experiment 1: N=36, Experiment 2: N=34). Green and orange points represent the mean RT of individual participants. a) and d) show mean RT individual scores in target absent trials when target-related audio sentences included a verb accompanied by a semantically incongruent visual clip vs. a semantically congruent visual clip for Experiment 1 and in Experiment 2, respectively. b) and e) show mean RT individual scores in target absent trials when target-related audio sentences included a noun accompanied by a semantically incongruent visual clip vs. a semantically congruent visual clip for Experiment 1 and in Experiment 2, respectively. c) and f) show mean RT total scores in incongruent trials for verb vs. noun (audio sentence with a target related verb vs. noun accompanied by a semantically incongruent visual clip) vs. congruent trials for Experiment 1 and in Experiment 2, respectively (audio sentence with a target related verb vs. noun accompanied by a semantically congruent visual clip).

